# The transcriptional regulator Sin3A balances IL-17A and Foxp3 expression in primary CD4 T cells

**DOI:** 10.1101/2022.04.19.488789

**Authors:** Laura Perucho, Laura Icardi, Elisabetta Di Simone, Veronica Basso, Amaia Vilas Zornoza, Teresa Lozano, Felipe Prosper, Juan José Lasarte, Anna Mondino

## Abstract

The Sin3 transcriptional regulator homolog A (Sin3A) is the core member of a multi-protein chromatin-modifying complex known to control gene transcription via epigenetic mechanisms. Its inactivation in developing thymocytes halts T cell maturation. We and others had previously shown that Sin3A controls STAT3 transcriptional activity. Given the role of STAT3 in the differentiation of T helper 17 cells critical in inflammatory disorders and against opportunistic infections, we asked whether Sin3A could also contribute to their differentiation. To this aim, we exploited CD4-Cre and CD4-CreER^T2^deleter strains for conditional and inducible Sin3A deletion in CD4 cell subsets. We report that Sin3A inactivation *in vivo* arrested thymocyte development at the double positive stage, hindering the characterization of mature T cells. At difference, tamoxifen-inducible Sin3A deletion proved permissive for *in vitro* proliferation of T cells in Th17 skewing conditions and the acquisition of memory markers. Transcriptional profiling indicated that while Sin3A inactivation imprinted T cells with a mTORC1 signaling gene signature, Sin3A deficient cells lacked the expression of IL-17A, the signature Th17 cytokine. This reflected a defective induction of *Il17a*, and also of the *Il23R* and *Il22* genes, which occurred in spite of proper upregulation of the lineage defining transcription factor RORγt. We found that Sin3A inactivation was paralleled by increased STAT3 phosphorylation and nuclear representation, and by higher fractions of IL-2 and FoxP3 expressing cells. Such events proved causally linked as inhibiting Foxp3 partially rescued IL-17A expression, and neutralizing IL-2 simultaneously lowered the representation of FoxP3^+^cells, while rescuing IL- 17A^+^ ones. Thus, together our data underline a previously unappreciated role for Sin3A in Th17 differentiation and the shaping of their immunoregulatory potential.

**Statement:** This study identifies a new role for the transcriptional regulator Sin3A in the shaping of Th17 cell differentiation. Data indicate that by controlling IL-2 expression, and mTORC1 signaling, it balances IL-17A and Foxp3 levels, shaping Th17 inflammatory potentials.

## Introduction

The Sin3 transcriptional regulator homolog A (Sin3A) is the core component of a multi-protein chromatin-modifying complex. While initially defined as a co-repressor, Sin3A is now considered a co-regulator, since it has co-repressor, co-activator and transcription factor properties (Chaubal & Pile, 2018). Sin3A contains four paired amphipathic helix domains (Ayer *et al*, 1995; Schreiber-Agus *et al*, 1995), each of them specific for particular transcriptional regulators. Through a class I histone deacetylases (HDAC) interaction domain (HID), Sin3A binds HDAC1 and HDAC2 and influences chromatin structures (Alland A, Muhle R, Hou Jr H, Potes J, Chin L, Schreiber-Angus N, 1997; Nagy *et al*, 1997; Laherty *et al*, 1997). Sin3A-associated HDAC1/2-activity is essential for embryonic stem cell survival (Fazzio *et al*, 2008), hematopoiesis, hematopoietic stem cell homeostasis and T and B lymphocyte development (Heideman *et al*, 2014). While HDAC activities are regarded as predominant, other chromatin modifiers were also found in Sin3A complexes, including histone methyltransferases (Nakamura *et al*, 2002), O-linked N-acetylglucosamine transferase (Yang *et al*, 2002), or members of the Swi/Snf nucleosome remodeling complex (Pal *et al*, 2003). Sin3A deletion caused the deregulation of genes involved in cell cycle, DNA replication, DNA repair, apoptosis, chromatin modification, mitochondrial metabolism and cell differentiation *in vitro* (Dannenberg *et al*, 2005), and early embryonic lethality (Dannenberg *et al*, 2005; Cowley *et al*, 2005; McDonel *et al*, 2012). Its conditional inactivation at the double negative (DN) stage of thymocyte development was found to cause a reduction in cellularity accompanied by an arrest in CD8 T cell selection and the appearance of dysfunctional CD4^+^ T cells (Cowley *et al*, 2005).

In previous studies, Sin3A was shown to regulate the transcriptional activity of the signal transducer and activator of transcription 3 (STAT3) (Gambi *et al*, 2019; Icardi *et al*, 2012), which is central to differentiation of Th17 cells. These cells mediate immunity against fungal infections and extracellular bacteria (Ye *et al*, 2001) through the secretion of several proinflammatory cytokines including IL-17A, IL-17F, IL-21, IL-22, IL-9, GM-CSF, TNFα and CCL20 (Dong, 2008; Liang *et al*, 2006). Complex signaling networks shape differentiation of Th17 cells (Yosef *et al*, 2013). Signals originated through the T-cell receptor (TCR) and co-stimulatory molecules, such as CD28, in the presence of IL-6 (Das *et al*, 2009) activate a number of transcription factors, among which is STAT3, which is required during early Th17 cell differentiation (Yang *et al*, 2007a; Zhou *et al*, 2007; Mathur *et al*, 2007). STAT3, in turn, controls the expression of the retinoic-acid-receptor related orphan receptor-γ (RORγt), the lineage-specific transcriptional regulator of Th17 (Ivanov *et al*, 2006). IL-6 and TGF-β also cause the STAT3-dependent expression of IL-21, which act via an autocrine loop to promote RORγt-dependent IL-17 expression (Korn *et al*, 2007; Wei *et al*, 2007; Korn *et al*, 2009). Transcription cofactors have also been shown to regulate Th17 cell development (Jiang *et al*, 2019). In addition to act on gene transcription, STAT3 has been shown to control mitochondrial Ca2^+^ and membrane potential, contributing to CD4^+^ T cell differentiation (Rincon & Pereira, 2018; Yang *et al*, 2015).

Given the role of STAT3 in Th17 cell differentiation, and the finding that Sin3A controls STAT3 in other cell types (Icardi *et al*, 2012; Gambi *et al*, 2019), we hypothesized that Sin3A could contribute to CD4^+^ T cell differentiation. To define the role of Sin3A and understand the consequences of its inactivation in CD4 cells, we crossed Sin3A^F/F^ mice with CD4-Cre, CD4-CreER^T2^ and Rosa26-CreER^T2^ deleter strains, allowing deletion of Sin3A *in vivo* at the DP stage of thymic development and *in vitro* in mature T cells. Data underline a previously unappreciated role for Sin3A CD4 T cells and in particular in fine tuning IL-2, mTOR and STAT3 signaling over the course of Th17 differentiation, balancing IL-17A and Foxp3 expression, ultimately shaping Th17 cell effector functions.

## Materials and methods

### Mice

Mice were housed and bred in the San Raffaele Institutional specific pathogen-free animal facilities. Mice used in this study were backcrossed with C57BL/6J mice (>10 generations). Mice carrying Sin3A-floxed alleles (Dannenberg *et al*, 2005) (Sin3A^F/F^; gently provided by Dr. Jan Tavernier, Ghent University and Hospital, Ghent University, 9000 Ghent, Belgium) were crossed with CD4-Cre (Lee *et al*, 2001) (gently provided by Dr. Paolo Dellabona, San Raffaele Institute), CD4-CreER^T2^ (gently provided by Burkhard Becher, University of Zurich), or Rosa26-CreER^T2^ (Ventura *et al*, 2007) (gently provided by Dr. Marco Bianchi, San Raffaele Institute) transgenic mice to generate Sin3A^F/F^ CD4-Cre^+/-^, Sin3A^F/F^ CD4-CreER^T2 +/-^ and Sin3A^F/F^ Rosa26-CreER^T2+/-^ mice, respectively. Sin3A^F/F^ mice were also crossed with OT II mice (gently provided by Dr. Andrea Annoni, San Raffaele Institute) and subsequently with Rosa26-CreER^T2+/-^ mice. Mice were screened by PCR. The following primers were used to detect the floxed (250 bp) or wild-type (150 bp) Sin3A allele: 5’- CAGATCCTATTCCAGGTGTCAAAG and 5’-CATGTTCATGTTTAGATATACTTCG. The presence (390 bp) of Cre was also investigated by PCR with the following primers 5’- CCTGGAAAATGCTTCTGTCCG and 5’-CAGGGTGTTATAAGCAATCCC. Studies were performed in accordance with the European Union guidelines and with the approval of the San Raffaele Institute Institutional Ethical Committee (Milan, Italy; IACUC-814).

### T Cell Activation and Differentiation

CD4^+^ T cells were isolated by negative selection from spleens and lymph nodes (Miltenyi Biotec, #130-104-454) according to manufacturer’s instructions (typically >95% purity). To initiate Sin3A deletion, CD4^+^ T cells were then cultured for 6 days in RPMI supplemented with 5% FBS, L-Glutamine, Pen/Strep, β-mercaptoethanol and IL-7 (5 ng/mL) in the absence or the presence of 4-OH tamoxifen (TAM, 1 µM). Cells (1 x 10^6^/mL) were then recovered and activated for 3 days on plate-bound anti-CD3 (BD, #553058) and soluble anti-CD28 (Biolegend, #102116) antibodies in Th17 or Treg skewing conditions. For Th17 differentiation, T cells were plated on 5 µg/ml plate-bound anti-CD3 and 5 µg/ml anti-CD28 antibodies in Iscove’s Modified Dulbecco’s Medium (IMDM) supplemented with 10% FBS, L-Glutamine, Pen/Strep, β-mercaptoethanol, 50 ng/mL IL-6, 1 ng/mL TGF-β, 10 ng/mL IL-1β, 10 µg/mL anti-IL-4 and 10 µg/mL anti-IFNγ. For Th0 differentiation, T cells were cultured on 1 µg/ml plate-bound anti-CD3 and 1 µg/ml soluble anti-CD28 antibodies in RPMI supplemented with 5% FBS, L-Glutamine, Pen/Strep, β-mercaptoethanol, 20 ng/mL IL-2 and 2.3 ng/mL TGF-β. For Th1 differentiation, cells were cultured as in Th0, supplementing cells with 3 µg/ml anti-IL-4 and 10 ng/ml IL-12. For Treg differentiation, T cells were cultured on 1 µg/ml plate-bound anti-CD3 and 5 µg/ml soluble anti-CD28 antibodies in RPMI supplemented with 5% FBS, L-Glutamine, Pen/Strep, β-mercaptoethanol, 20 ng/mL IL-2 and 2.3 ng/mL TGF-β. All cytokines used were purchased from Peprotech. After 3 days of activation, cells were stimulated with Phorbol 12-myristate 13-acetate (PMA) (10 ng/mL, Sigma, #P8139) / ionomycin (1 µg/mL, Sigma, # I0634) for 4 h, adding Brefeldin A (BFA) (5 µg/mL, sigma, # B7651) for the last 2h of culture. As control, half of the cells were treated only with BFA. In some experiments, anti-IL-2 blocking antibody (1 µg/ml unless specified, BioCell #BE0043-1) or the Foxp3 inhibitor P60(Casares *et al*, 2010) (20 µM) were provided during the 3 days Th17 differentiating cultures. In experiments using OTII^+/-^ Sin3A CreERT2 mice, CD4 T cells were purified as above, kept for 6 days in RPMI supplemented with 5% FBS, L-Glutamine, Pen/Strep, β-mercaptoethanol and IL-7 (5 ng/mL) in the absence or the presence of 4-OH tamoxifen (TAM, 1 µM) and then cultured for 3 days in Th17 conditions in the presence of 1.25 µM ovalbumin (OVA).

### Flow cytometry

Cells were collected, washed in PBS and stained with a viable dye (Zombie Aqua, Biolegend, #423101) for 20 min on ice in the dark. Cells were then resuspended in Fc-blocking solution (anti CD16/CD32 antibody 2.4G2, homemade), incubated for 5 minutes and then labeled with indicated antibodies for 15 min at 4°C in the dark. Cells were then washed and fixed for 30 min to ON at 4°C (Foxp3/Transcription Staining Buffer Set, eBioscience, #00-5523-00). Fixed cells were permeabilized using the same buffer set and intracellular stainings were performed for 30 min at 4°C in the dark. Stained cells were collected and measured on a BD FACSCanto3 machine. Analyses were performed using the FlowJo software. The list of antibodies adopted in the study is provided Supplementary Table S2).

### Apoptosis and Cell Proliferation

To measure apoptosis, surface levels of Annexin V were investigated by flow cytometry. 1×10^5^ cells were washed and resuspended in 100 µl of ice-cold binding buffer (140 mM NaCl, 4 mM KCl, 0.75 mM MgCl2 and 10 mM HEPES). 10 min before acquisition, anti-AnnexinV-FITC antibody (5 µl, BD, #51-65874X), 7-AAD (5 µl, BD, #559925) and 1.75 µl of 85 mM CaCl_2_ were added. Proliferation was determined by labeling T cells with the Tag-IT Violet Proliferation Cell Tracking Dye (Biolegend, #425101), according to manufacturer instructions. Briefly, cells were washed twice with PBS, resuspended in PBS (100 x 10^6^ cell/mL) and incubated with Tag-It Violet Proliferation Cell Tracking Dye (5 µM) for 20 min RT in the dark. Then, an equal volume of FBS was added to the sample, and cells were finally washed with and 10 ml cell culture medium.

### Cell cycle analysis

CD4^+^ T cells (1×10^6^) were washed twice with cold PBS, resuspended in 500 µl PBS, fixed by adding 4.5 ml ice-cold 70% ethanol while vortexing and incubated for at least 2 hours at 4°C. Samples were centrifuged, washed with PBS for 15 min RT, incubated in 300 µl of 1 µg/ml DAPI 0.1% Triton X-100 in PBS for 30 min and acquired in linear mode at low speed.

### RNA-Seq analysis

Illumina strand specific mRNA Roughly 100ng of high-quality total RNA (RIN >8) was used for the transcriptomic interrogation of control and SIN3A deleted CD4 T cells using Illumina’s Stranded mRNA Prep ligation according to the manufacturer’s instructions. Briefly, Oligo(dT) magnetic beads were used to purify and capture the mRNA molecules in the total RNA. The purified mRNA was fragmented and copied into first strand complementary DNA (cDNA) using reverse transcriptase and random primers. A second strand cDNA synthesis step removed the RNA template while incorporating dUTP in place of dTTP in order to preserve strand specificity. Next, double-stranded cDNA was A-tailed, then ligated to Illumina anchors bearing T-overhangs. PCR-amplification of the library allowed the barcoding of the samples with 10bp dual indexes and the completion of Illumina sequences for cluster generation. Libraries were quantified with Qubit dsDNA HS Assay Kit and their profile was examined using Agilent’s HS D1000 ScreenTape Assay. Sequencing was carried out in an Illumina NextSeq2000 using paired end, dual-index sequencing (Rd1: 59 cycles; i7: 10 cycles; i5: 10 cycles, Rd2 59 cycles) at a depth of 30 million reads per sample.

RNA-seq reads are trimmed using Trim Galore v0.4.4 using default parameters to remove the Nextera adapter sequence. Mapping is performed using STAR (2.6) against the mouse NCBIM37 genome, guided by gene models from Ensembl annotation release 68. Quantification and generation of gene expression matrices were performed with the function featureCounts, implemented in the R package Rsubread. Aligned fragments are imported into RStudio and before statistical analysis, the function filterbyExpr, implemented in the R package edgeR, was used to determine genes with enough counts for further analyses. Differential gene expression analysis is performed using the DESeq2 algorithm within R and RStudio. Gene set enrichment analysis was carried out using GSEA software (https://www.gsea-msigdb.org/). The data discussed in this publication have been deposited in NCBI’s Gene Expression Omnibus (Edgar *et al*, 2002) and will be accessible through GEO Series accession number GSE_196615_ (https://www.ncbi.nlm.nih.gov/geo/query/acc.cgi?acc=GSE196615).

### RT-PCR and Real-time PCR

Total RNA was purified using ReliaPrep RNA Miniprep System (Promega, #Z6011), according to manufacturer’s instructions. RNA was retrotranscribed for 1 hour at 42°C with retro transcriptase M-MLV (Invitrogen, #28025-013) in the presence of oligo(dT)_15_ Primer (Promega, #C1101), dNTPs (Euroclone), and RNAsin Ribonuclease Inhibitor (Promega, # N2111). The cDNA was diluted 10 times to perform qRT-PCR with SYBR Green Master Mix (Applied Biosystems, #4309155). The list of primers adopted in the study is provided in Suppl Table S1). *C*_t_ values were normalized to housekeeping genes expression by the ΔΔ*C*_t_ method.

### Western Blot

Total lysates were recovered in 2% SDS Tris-HCl 65 mmol/L pH 6.8 lysis buffer and sonicated. Whole cell, cytosolic and nuclear extracts were obtained using the REAP protocol(Suzuki *et al*, 2010). Lysates corresponding to the same number of cells were separated by SDS-PAGE, transferred to nitrocellulose membrane, blocked with phosphate buffered solution (PBS)-5% milk, and subjected to immunoblotting. The following antibodies were used: Anti-STAT3 (Cell Signaling, #9139), anti-(Ser727) STAT3 (Cell Signaling, #9134), anti-β-ACTIN (Cell Signaling, #3700), anti-SIN3A (Cell Signaling, #7691). Quantification of protein levels was performed by measuring the mean grey value of the bands using ImageJ.

### Statistical Analyses

Unless otherwise indicated, a paired two-tailed student’s t test was used. P values < 0.05 were considered significant. Error bars represents standard deviations (SD). GraphPad Prism 9 was used to perform statistical analyses.

### Logistic model

A logistic model was adopted to assign the distribution of TAM-treated or untreated cells based on selected markers. The R stats package (version 4.0.2) for logistic model fitting and the R ggplot2 package (version 3.3.3) were used and run in Rstudio (version 1.0.153).

## Results

### CD4-Cre-driven Sin3A inactivation halts proper thymocyte development

A previous report found that Sin3A deletion in DN thymocytes via Lck-driven Cre expression hindered CD8 thymocyte development and caused post-thymic accumulation of dysfunctional CD4^+^ T cells (Cowley *et al*, 2005). To better understand the contribution of Sin3A to CD4^+^ T cell development and differentiation, we crossed Sin3A^F/F^ with CD4-Cre mice (Sin3A-CD4Δ) and CD4-CreER^T2^ deleter strains, which allows evaluating the impact of Sin3A inactivation beyond the DN stage of thymocyte development, i.e. over the course of DP to SP transition. Flow cytometry (FACS) analyses revealed that both CD4SP and CD8SP cells were significantly underrepresented in Sin3A-CD4Δ mice when compared to controls (Fig 1A-B). This correlated with Sin3A expression, comparable in DN cells and respectively reduced and completely lost in DP and SP cells derived from Sin3A-CD4Δ mice compared to controls (Fig 1C). Impaired thymic development also caused a severe reduction in the frequency of CD4^+^ and CD8^+^ T cells in secondary lymphoid organs (Fig 1D-E). Of note, remaining CD4^+^ T cells in the spleen of Sin3A-CD4Δ mice had acquired a CD44^high^ CD62L^low^ effector memory phenotype (Fig 1F-H). This was likely the results of cells having escaped CD4-Cre-driven Sin3A deletion (Fig 1H), and having acquired a memory phenotype as a result of lymphopenia induced proliferation (Sckisel *et al*, 2017). Similar results were obtained when administering tamoxifen to Sin3A^F/F^CD4-CreER^T2^ mice. Tamoxifen administration caused a progressive loss of circulating mature CD4^+^ T cells secondary to a developmental arrest (Supp. Fig. 1). Thus, conditional deletion of Sin3A at the DP stage unequivocally defines a role for Sin3A in thymocyte development and underlines the need for post-thymic inactivation for the analysis in mature T cells.

**Figure 1.**
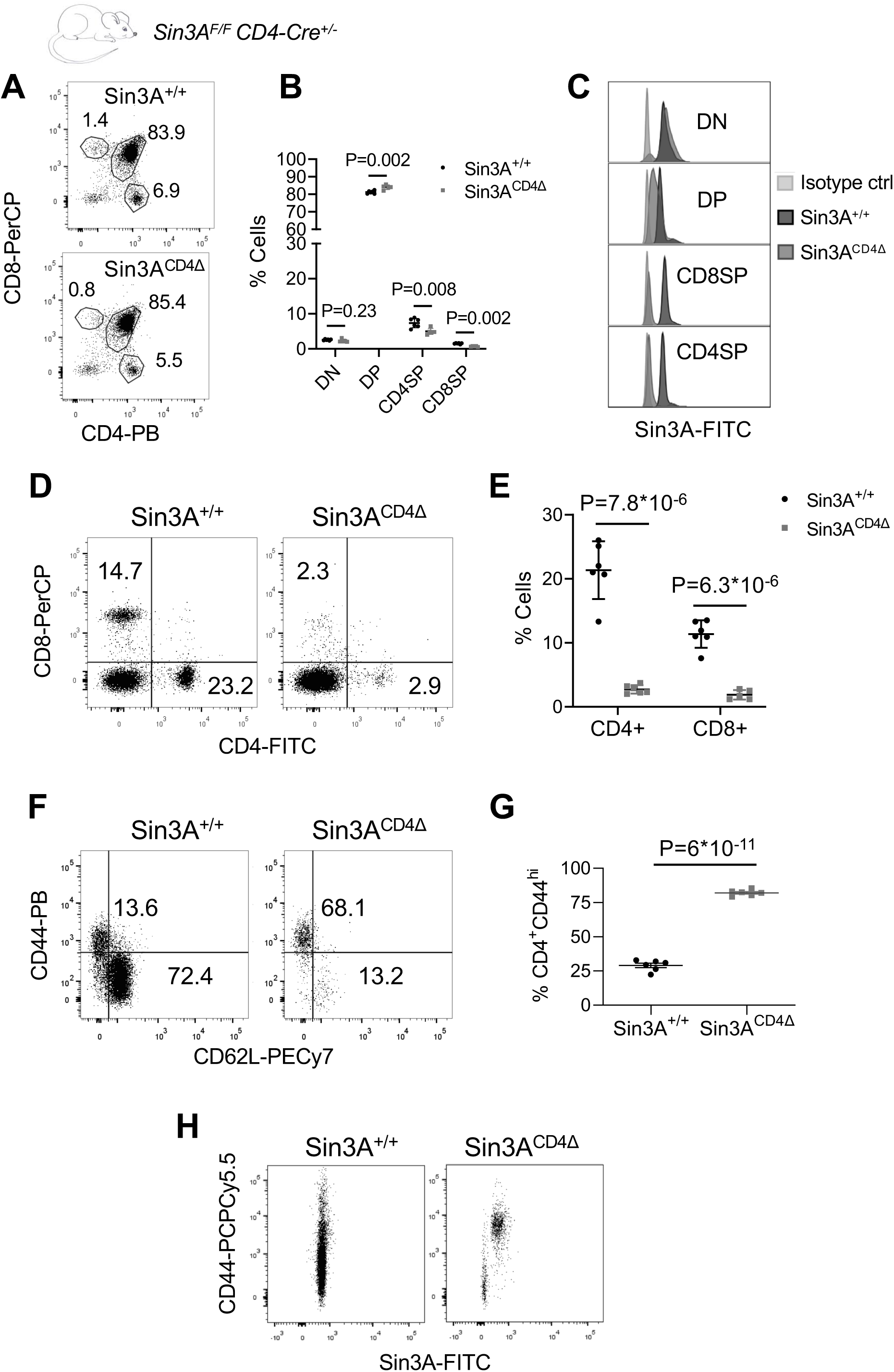
CD4-cre-driven Sin3A deletion halts further CD4 and CD8 thymocyte development. **A-B**. Representative flow cytometry analysis of CD4^+^ and CD8^+^ thymocytes (**A**), and frequencies of DN, DP and SP cells in Sin3A^+/+^ and Sin3A^CD4Δ^ mice (**B**). N=5. **C**. Sin3A intracellular levels in DN, DP, CD8SP and CD4SP cells from Sin3A^+/+^ and Sin3A^CD4Δ^ mice. **D-E**. Representative FACS of splenic CD4^+^ and CD8^+^ cells (**D**) and frequencies (**E**). N=6. **F-G**. Representative dot plots of CD44 and CD62L levels of gated CD4^+^ viable T cells (**F**) and quantification (**G**). N=6. **H**. Representative dot plots of CD44 and Sin3A levels in Sin3A^+/+^ and Sin3A^CD4Δ^ CD4^+^ T cells. **B, E, G.** Bars: Mean. Error bars: SD. Two-tailed unpaired Student’s T-test.

### Sin3A deletion in mature CD4 T cells imprints a mTORC1 signature and hinders Th17 cell differentiation

We first investigated Sin3A expression in purified CD4^+^ T cells, and found it to be upregulated by day 3 of anti-CD3/CD28 activation in various skewing condition (Fig. 2A) and over the course of Th17 differentiation (Fig. 2B). Next, we cultured purified CD4^+^ T cells obtained from CD4-Cre CreER^T2 +/-^ Sin3A^F/F^ or Rosa26-CreER^T2 +/-^ Sin3A^F/F^ mice for 6 days in interleukin 7 (IL-7) in the absence or the presence of 4-hydroxytamoxifen (TAM). This was adopted to allow Cre-mediated Sin3A deletion in mature T cells while avoiding activation (Tan *et al*, 2001). To investigate the contribution of Sin3A to Th17 differentiation, cells were then cultured for an additional 3 days in Th17 skewing conditions (refer to Fig. 2C for a schematic experimental outline). Western blot and FACS analysis verified the loss of Sin3A expression in TAM-treated cells (Fig. 2D and E). Such effect was more robust and consistent in T cells derived from Rosa26-CreER^T2 +/-^ Sin3A^F/F^ mice (not shown), which were then adopted for all further studies. Over the course of Th17 skewing, both control and TAM-treated cells completed several rounds of cell division, indicated by the dilution of the CFSE vital dye (Fig 3A-B). Fewer TAM-treated cells completed more than 3 cell division cycles (Fig. 3B). This was paralleled by lower cellular yields (Fig 3C), and a slightly increase in apoptotic cells (Supp. Fig 2A-B), and in cells in the S phase of the cell cycle (Supp. Fig 2C). To some extent, this recapitulates the phenotype of mouse embryonic fibroblasts, embryonic stem cells (Fazzio *et al*, 2008) and pluripotent cells (Dannenberg *et al*, 2005).

**Figure 2.**
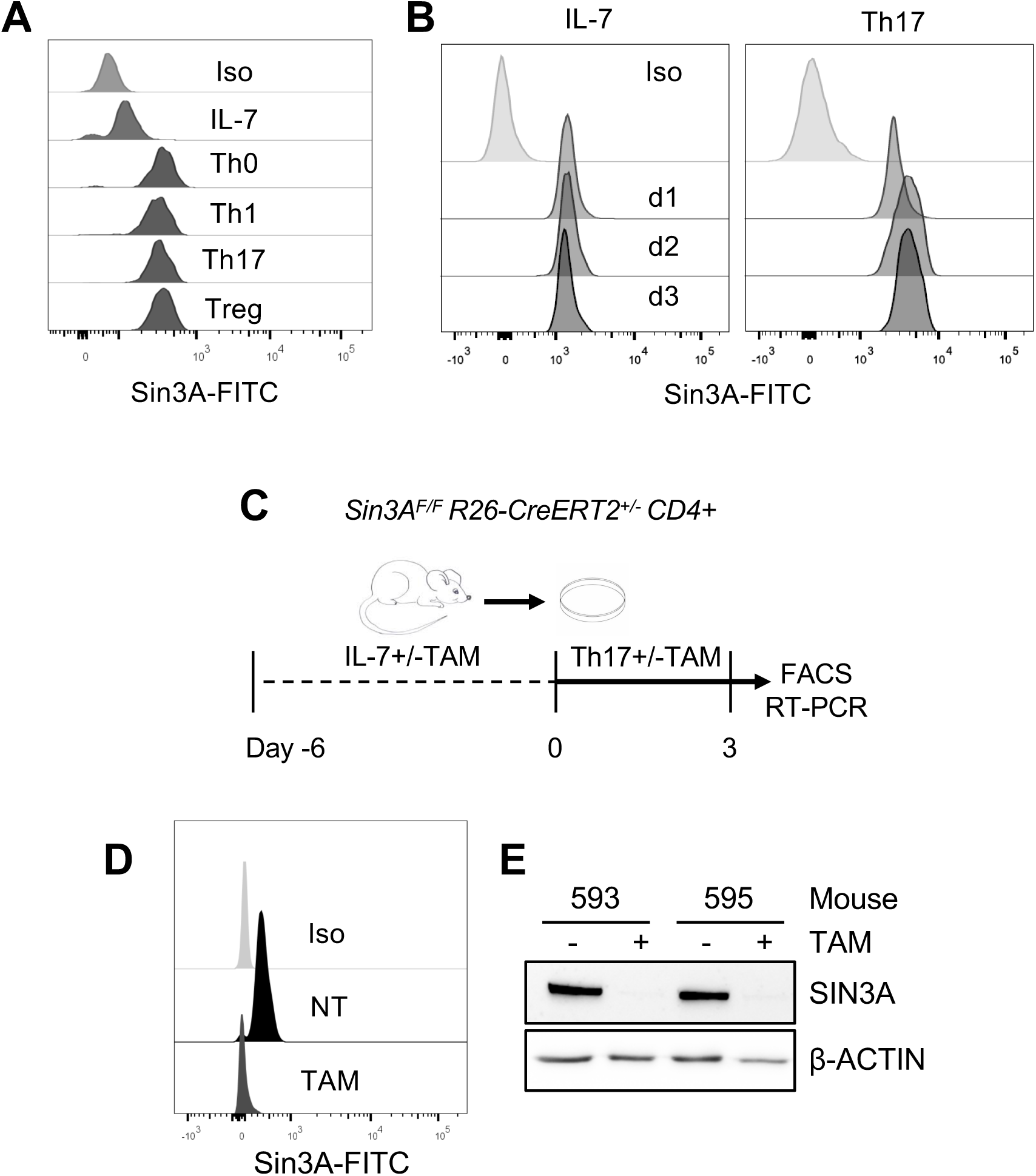
Sin3A levels are increased over the course of activation and differentiation. **A.** Representative flow cytometric analysis of Sin3A expression in wild-type CD4^+^ T cells kept for 3 days in IL-7 or differentiated to Th0, Th1, Th17 and Treg. **B**. Kinetics of Sin3A upregulation in Th17 conditions. **C**. Graphic representation of *ex-vivo* deletion of Sin3A. CD4^+^ T cells were purified from secondary lymphoid organs derived from Sin3A^F/F^ Rosa26-CreER^T2+/-^ mice, and kept in IL-7 for 3 days in in the absence or the presence of 4-hydroxy tamoxifen (TAM). Cells were then harvested and cultured for an additional 3 days in Th17 skewing conditions in the absence or the presence of TAM. **D.** Flow cytometric and **E.** Western blot analyses of Sin3A levels.

**Figure 3.**
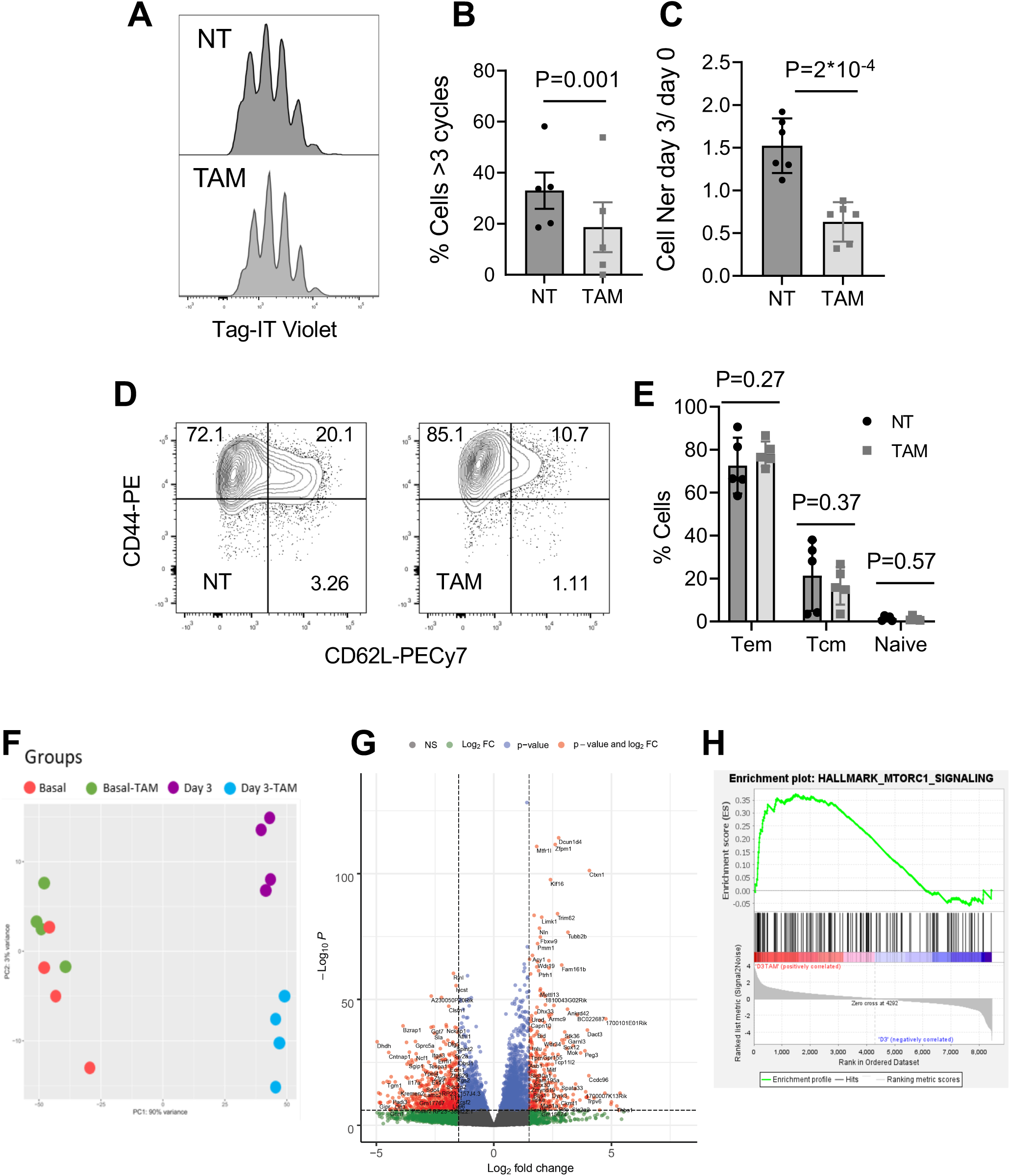
Conditional deletion of Sin3A allows cell differentiation and imprints T cells with a mTORC1 signature. Sin3A^F/F^ Rosa26-CreER^T2^ CD4^+^ T cells were treated as described in Figure 2. **A.** Proliferation profiles of untreated (NT) and TAM-treated (TAM) cells labeled with Tag-IT Violet, and kept for 3 days in Th17 conditions. **B**. Quantification of cell division cycle according to Tag-IT Violet dilution profiles, N=5. **C**. Relative cell numbers (day 3/day 0). N=6. **D.** Contour plots depicting relative CD44 and CD62L expression after gating on viable lymphocytes **E.** Bar plots showing relative frequencies of effector memory (Tem), central memory (Tcm) and naïve cells viable lymphocytes, N=5. **F-H**. RNA-seq transcriptomic analysis of CD4^+^ T cells derived from Rosa26-CreER^T2^ Sin3A^F/F^ mice cultured for 3 days in IL-7 in the absence or the presence of tamoxifen (basal, basal TAM) and then cultured for 3 additional days in Th17 polarizing conditions (Day3 or Day3TAM). Principal component analyses (**F**). Volcano plot of differentially expressed genes (FDR<0.05; vertical dashed lines indicate fold change values Long2 fold change 1-1) (**G**) Gene Set Enrichment analysis (**H**) highlights a signature compatible with an MTORC1 signaling pathway (Genes up-regulated through activation of mTORC1 complex).

In spite of the mild proliferative defect, Sin3A deficient cells acquired a typical memory phenotype in response to Th17 skewing conditions. Indeed, while IL-7-cultured CD4^+^ T cells retained a CD44^low^, CD62L^high^ naïve phenotype, and lacked expression of activation markers such as CD25, PD-1 or CD69 (Supp. Fig 3A), most of Sin3A suffidient and deficient Th17- skewed T cells acquired a CD44^high^, CD62L^low^ effector memory phenotype (Fig 3E), and upregulated CD69, CD25, PD-1 (Supp Fig 3B-C).

To investigate T cell behavior in response to a nominal antigen, we generated ovalbumin-specific TCR transgenic OTII^+/-^ Sin3A^F/F^ CreERT2^+/-^ mice. CD4 T cells purified from these mice were then compared to those derived from OTII^+/-^ Sin3A^+/F^ CreERT2^+/-^ littermates to also control for Cre-mediated toxicity (Loonstra A, Vooijs M, Beverloo HB, Al Allak B, van Drunen E, Kanaar R, Berns A, 2001). CD4 T cells were purified and treated or not with TAM, labeled with CFSE and then stimulated with ovalbumin. Data depicted in Supp. Fig 4A-B indicate that antigen stumulation induced proliferation of both Sin3A sufficient and deficient cells. Again, although OTII cells lacking Sin3A (OTII^+/-^ Sin3A^+/F^ CreERT2^+/-^) revealed a mild proliferative and survival defect compared to controls, they similarly acquired a CD44^high^, CD25^high^ phenotype (Supp. Fig. 4C). Thus, together these data indicate that deletion of Sin3A in mature T cells is compatible with TCR-driven proliferation and differentiation into memory cells.

To obtain an unbiased overview of the impact of Sin3A inactivation on Th17 differentiation, we next performed RNA-seq analysis (Fig 3F-H). To this aim, highly purified CD4^+^ T cells were kept for 6 days in IL-7 without or with TAM (Basal or BasalTAM), and then cultured for 3 additional days in Th17 polarizing conditions (Day3 or Day3TAM) (Fig 3F-H). RNA was then isolated and sequenced for transcripts differential expression. Most of the transcript variation segregated CD4^+^ T cells by the Th17 polarizing conditions. First principal components (PC1) analyses accounted for 90% of the variance between Th17 polarized T cells and unstimulated cells (basal conditions), and clearly differentiated Day3 samples (Sin3A sufficient) from Day3TAM ones (Sin3A deficient) (Fig. 3 F). Comparison between Day3TAM and Day3 T cells yielded a total of 9386 differentially expressed genes (p<0.05). Among them, 1139 genes were upregulated and 2282 were downregulated in Day3TAM T cells compared to controls (p<0.05), when considering genes with a fold change >1, as represented in the Volcano plot (Fig. 3G). Gene set enrichment analysis (GSEA) was next performed comparing Day3TAM and Day3 samples. We found a significant enrichment of genes implicated in mTORC1 signaling pathway (Genes up-regulated through activation of mTORC1 complex) (Fig. 3 H, Nominal p Value < 0.001, FDR q-value: 0.147). The finding that Sin3A inactivation imprints memory T cells with an active mTORC1 signaling signature, is strongly suggestive of a role of Sin3A in Th17 differentiation, given the notion that mTORC1 controls STAT3 phosphorylation, and Th17 differentiation.

Thus, we next investigated the expression of IL-17A, the signature cytokine for Th17 cells. We found that while a fraction of control cells expressed IL-17A after PMA and Ionomycin stimulation (PI), TAM-treated Sin3A-deficient cells mostly failed to do so (Fig. 4A). At difference, cells produced IFNγ to comparable extents (Fig. 4B). Defective IL-17A could not be attributed to differences in proliferation, as it was observed among Sin3A sufficient and deficient cells having undergone similar rounds of cell division (Suppl. Fig.5A-C). Of note, a fraction of Sin3A deficient cells preserved IL-17A expression (Suppl. Fig.5B).

**Figure 4.**
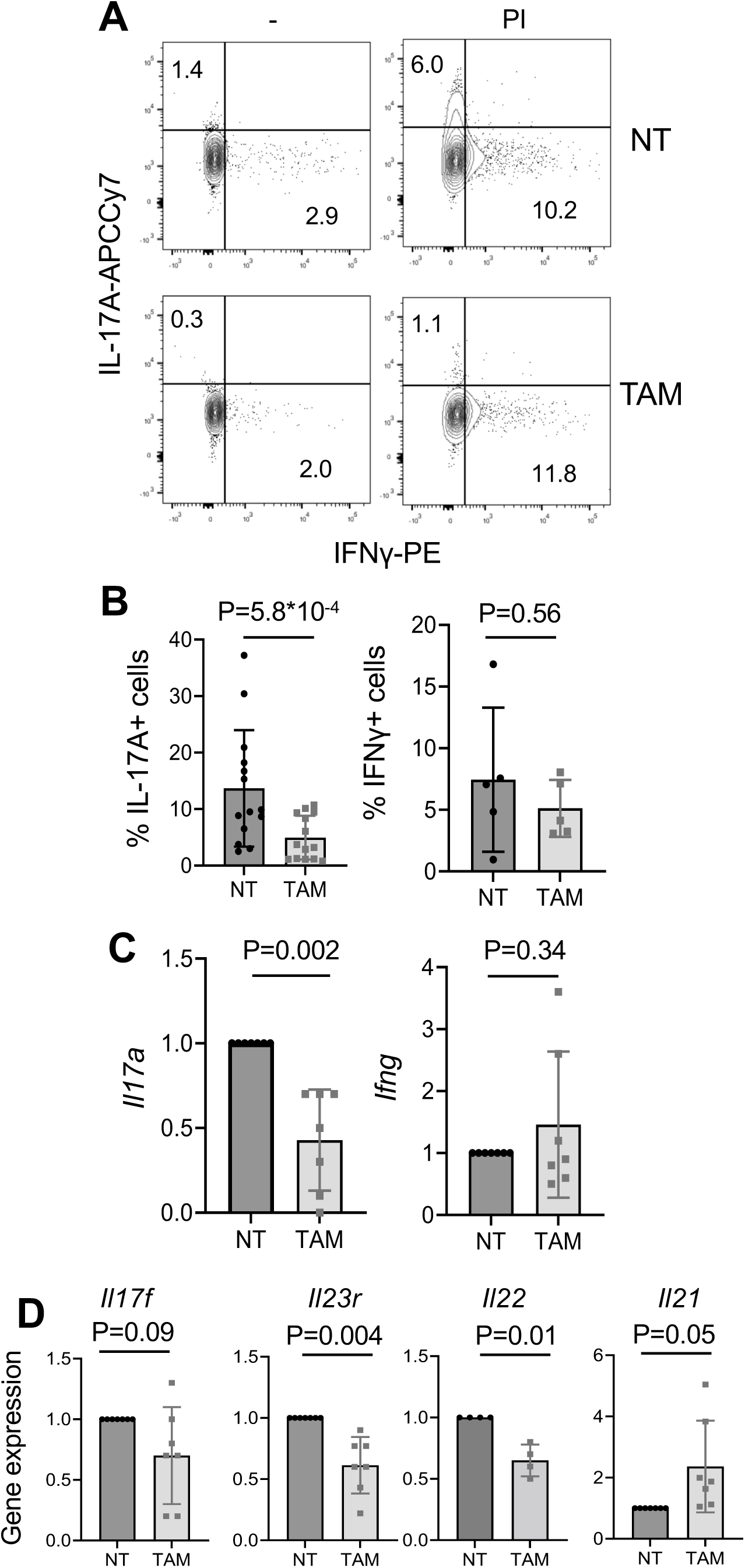
Sin3A deficient T cells fail to differentiate in Th17 skewing conditions. Sin3A^F/F^ Rosa26-CreER^T2^ CD4 T cells treated or not with TAM were kept for 3 days in Th17-skewing conditions as described in Figure 2. Cells were then left untreated or stimulated with PMA/ionomycin (PI) for 4h, and analyzed by intracellular cytokine staining (**A-B**) or RT-PCR (**C-D**). **A.** Representative dot plots and **B.** bar plots showing the percentage of cells expressing IL-17A^+^ and IFNγ^+^ (N=5). **C-D**. RT-PCR analysis of indicated gene expression after PI stimulation. Relative levels are depicted: *IL17a, ifng*, *Il17f, Il21* (N=7), *Il22* (N=4). *Il23r* levels were analyzed without re-stimulation (N=7). Bars: Mean. Error bars: SD. Two-tailed paired Student’s T-test.

**Figure 5.**
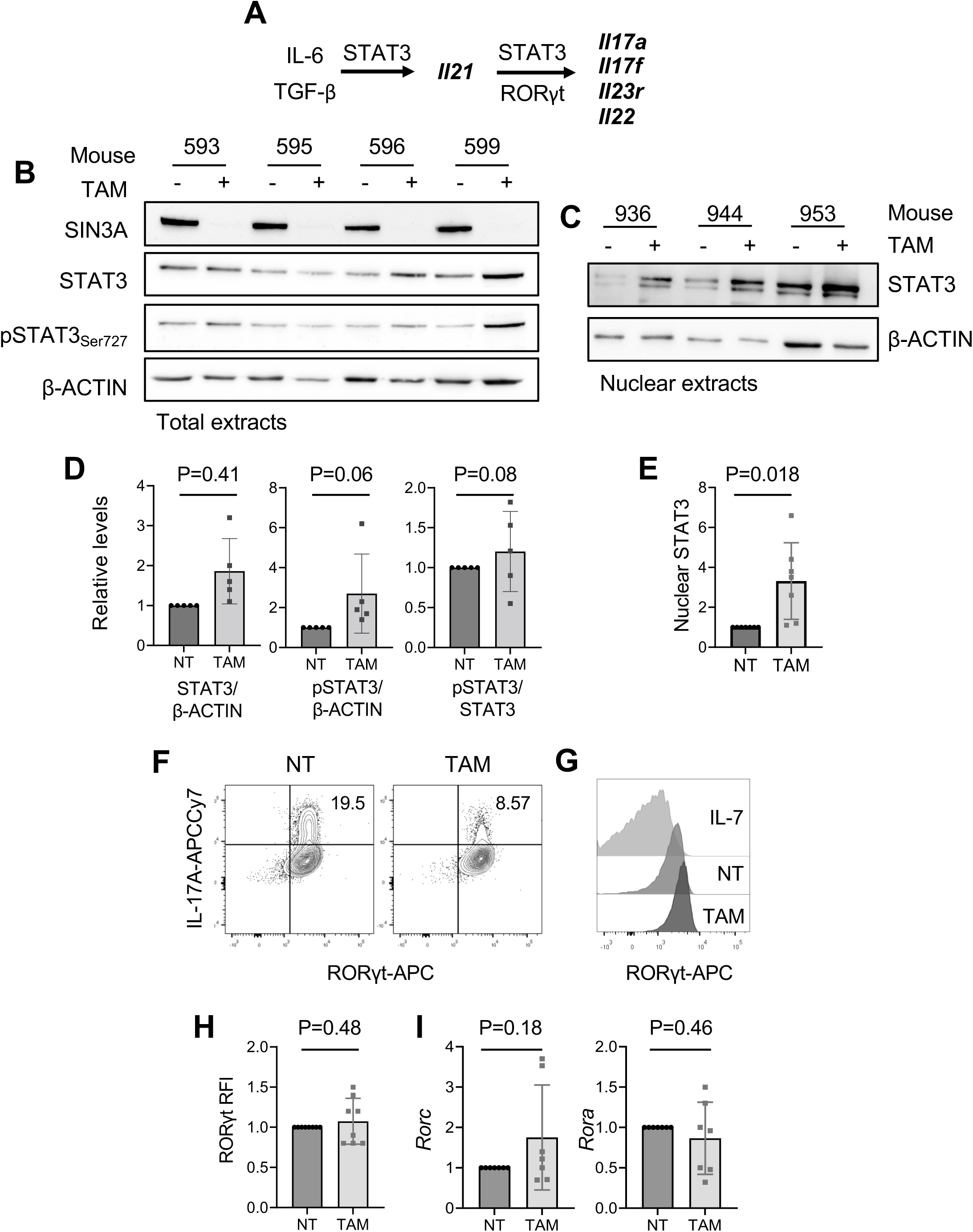
STAT3 and RORγt are expressed in Sin3A-deficient cells lacking IL17A. **A**. Schematic representation of STAT3 and RORγt-driven events over Th17 differentiation. **B-E.** Representative western blot of total (**B**) and nuclear (**C**) STAT3 and pSTAT3 (ser727) in control (NT) and Sin3A-deleted Th17-skewed CD4^+^ T cells (TAM), and quantification of relative representation over independent experiments (**D-E**), N=5. Bars: Mean. Error bars: SD.Two-tailed paired Student’s T-test, except in C. pSTAT3/β-ACTIN: Wilcoxon matched-pairs signed rank test (not normally distributed according to Shapiro-Wilk test). **F**. Contour plots depicting IL-17A and RORγt levels in viable PMA and Ionomycin stimulated control (NT) and TAM-treated CD4+ T cells. **G.** Representative histograms of RORγt protein levels and **H.** relative expression (Relative Fluorescence Intensity, TAM relative to NT), N=8. **I.** RT-PCR analysis of *Rorc* (N=6) and *Rora* (N=7) gene levels. Relative expression is shown.

RT-PCR analysis pointed towards a transcriptional defect, as Sin3AΔ cells revealed lower *Il17a* RNA levels compared to controls (Fig 4C), while *Ifng* levels were similar (Fig 4C-D). In addition to *Il17a*, also the expression of *Il17f, Il23r* and *Il22* were lower in Sin3A deleted cells, while *Il21* was significantly increased when compared to controls (Fig. 4D). Thus, post-thymic genetic inactivation of Sin3A hinders proper differentiation of Th17 cells.

### STAT3 and Foxp3 upregulation reflect defective *Il17a* expression in RORγt^+^ Sin3A-deficient CD4^+^ T cells

Differentiation of Th17 helper cells depends on signals from IL-6, in combination with TGF-β, IL-23 and IL-21. These signals lead to the activation of STAT3 and the expression of RORγt, the Th17 lineage defining transcription factor. RORγt then acts in synergy with STAT3 (Ciofani *et al*, 2012), to promote the production of IL-17A, IL-17F, and IL-22 (Ivanov *et al*, 2006; Zhou *et al*, 2007; Manel *et al*, 2008) (Fig 5A).

As Sin3A controls STAT3 transcriptional activity in several cell types (Icardi *et al*, 2012; Gambi *et al*, 2019), we first investigated relative STAT3 expression, phosphorylation and nuclear representation. We found a trend towards increased levels of total STAT3 and of its phosphorylation on Ser727 in cells lacking Sin3A compared to controls (Fig 5B, D). In addition, nuclear STAT3 levels were significantly higher in Sin3A-deficient cells compared to controls (Fig 5C, E). This is in line with the transcriptomic data suggesting an upregulation of mTORC1 (Fig. 3) which is upstream of STAT3, and suggest that STAT3 might be more transcriptionally active in cells lacking Sin3A, which is corroborated by higher *Il21* levels (Fig. 4D).

Next, we analyzed RORγt regulation. Data indicated RORγt to be expressed at comparable levels in Th17 skewed Sin3A sufficient and deficient cells both at the protein (Fig 5F-H) and mRNA level (Fig. 5I). Thus, cells lacking Sin3A preserve expression of STAT3 and RORγt. However, while *Il21*, which mainly relies on STAT3 (Wei *et al*, 2007) was expressed at higher levels in cells lacking Sin3A (Fig. 4D), *Il17a*, *Il23r*, and *Il22*, which are concomitantly regulated by STAT3 and RORγt (Ivanov *et al*, 2006; Durant *et al*, 2011; Ciofani *et al*, 2012), failed to be properly induced. These data support the notion that STAT3 and not RORγt reveal transcriptional active in Th17 cells lacking Sin3A.

STAT3 also controls Foxp3 expression (Pallandre *et al*, 2007). In turns, Foxp3 can directly interact with RORγt, and inhibit *Il17a* transcription (Ichiyama *et al*, 2008; Zhou *et al*, 2008). We thus asked whether the absence of Sin3A, could favor the expression of Foxp3 in spite of Th17 skewing conditions, and this might account for defective of *il17a* upregulation. Data reported in Figure 6 indicate that cells lacking Sin3A were indeed enriched for Foxp3^+^ cells (Fig 6A-B), likely due to increased *Foxp3* mRNA levels (Fig. 6C). Of note, when analyzing individual cells, Foxp3 and IL-17A proteins were mutually exclusive (Fig. 6A), supporting the possibility that gained Foxp3 expression could be causally linked to inhibition of *Il17a*, and the absence of IL-17A.

**Figure 6.**
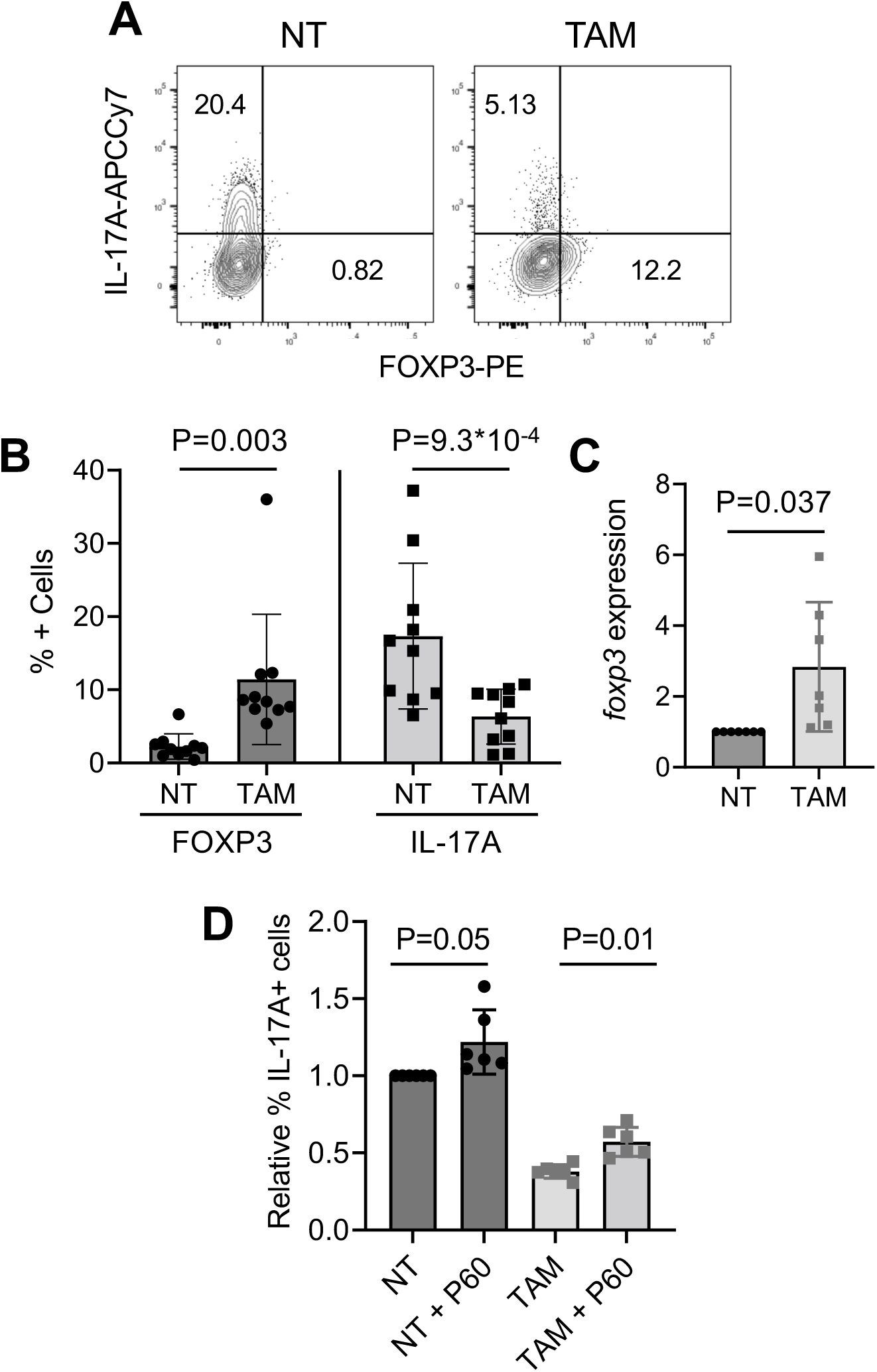
Sin3A inactivation leads to the upregulation of Foxp3, able to limit IL-17A. Cells were cultured as depicted in Figure 2 and analyzed after PMA and Ionomycin stimulation for 4h. **A.** Representative contour plots and **B.** percentages of IL-17A^+^ and Foxp3^+^ cells, N=10. **C**. RT-PCR analysis of *Foxp3* expression N=7. **D**. Cells were cultured as depicted in Figure 2 in the absence or the presence of the P60 Foxp3 inhibitor. Cells were then harvested and stimulated with PMA and ionomycin for 4h. Bar plots report the percentage of IL-17A+ cells relative to untreated controls, N=6. Error bars: SD. Two-tailed paired Student’s T-test.

We thus asked whether interfering with FoxP3 activity would rescue IL-17A expression. To this aim, we took advantage of P60, a cell permeable 15-mer synthetic peptide reported to bind to Foxp3 and to prevent its nuclear translocation (Casares *et al*, 2010). Sin3A sufficient and deficient cells were cultured in Th17 skewing conditions in the absence or the presence of P60, and then analyzed for IL-17A expression. We found that adding the P60 inhibitor over the course of Th17 differentiation significantly increased the percentage of IL-17A^+^ cells (Fig. 6D), with no effects on cell viability (Supplementary Fig. 6A). Although this was the case for both Sin3A sufficient and deficient cells, the rescue was more statistically relevant in the latter. Thus, gained Foxp3 expression in cells lacking Sin3A contributes at least in part, to reduced IL-17A levels.

### Sin3A deletion favors IL-2 expression, which is causally linked to Foxp3 upregulation and IL-17A inhibition

IL-2 promotes mTORC1 signaling (Ray *et al*, 2015), STAT3 and STAT5 phosphorylation and activation and Foxp3 expression and stability (Chen *et al*, 2011; Setoguchi *et al*, 2005; Zorn *et al*, 2006). As Sin3A was previously reported to negatively regulate *Il2* transcription by bringing HDAC to its promoter (Lam *et al*, 2015), we reasoned that the loss of Sin3A could unleash increased IL-2 expression, and this could contribute to the upregulation of FoxP3 and defective *Il17a* expression. By FACS analysis, we found that Sin3A deficient cultures were indeed enriched in IL-2 producing cells (Fig 7A), which was corroborated by higher *Il2* mRNA levels (Fig 7C), and a rise in phosphorylated STAT5, and p70S6 kinase (Suppl. Fig. 7). These data provide a mechanistic explanation for the finding that Sin3A-deficient cells were characterized by the mTORC1 signaling signature. Thus, Sin3A inactivation unleash *Il2* transcription in Th17 conditions, and this might cause il17a/IL-17A inhibition

**Figure 7.**
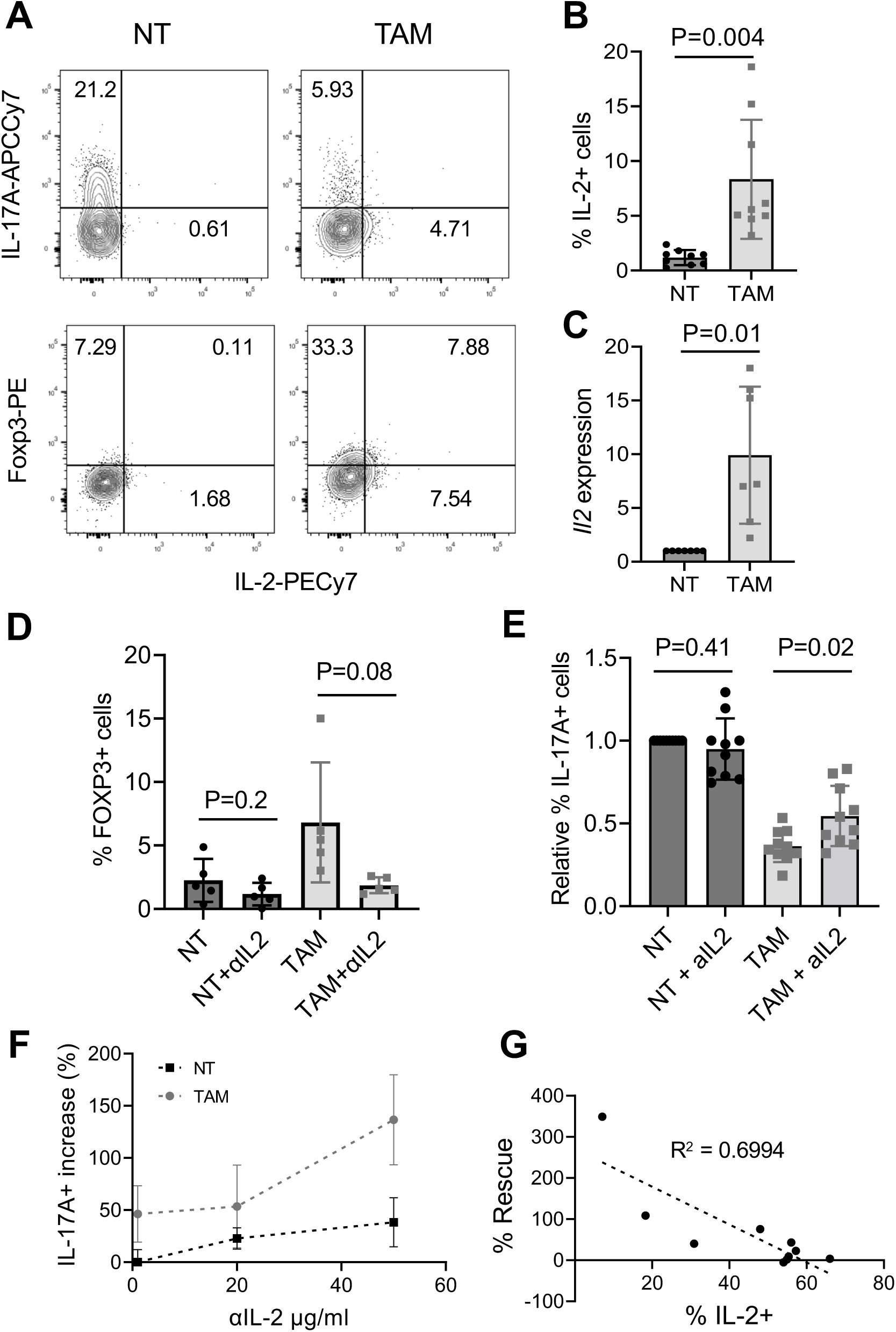
Sin3A inactivation favors IL-2 upregulation, which neutralization halts Foxp3 and rescues IL-17A expression. **A-B)** Cells were cultured as depicted in Figure 2 and then stimulated with PMA and Ionomycin for 4h. **A**. Representative contour plots and **B.** percentages of IL-2^+^ cells, N=7. **C.** RT-PCR analysis of *Il2* expression, N=7. **D-E**) Cells were cultured as depicted in Figure 2 in the absence or the presence of a neutralizing anti-IL-2 mAb. **D.** Bar plot depicting the frequency of Foxp3^+^, N=5 and **E.** relative IL-17A^+^ cells, N=10. **F.** Relative increase in IL-17A^+^ Sin3A sufficient and deficient cells with respect to the dose of anti-IL-2 mAb, N=3. **G**. Correlation of the relative increase in IL-17A^+^ cells in the presence of anti-IL-2 antibody and the percentage of IL-2^+^ cells in TAM-treated cells. Bars: Mean. Error bars: SD. Two-tailed paired Student’s T-test.

To directly challenge the role of IL-2, we investigated whether neutralizing IL-2 during Th17 skewing would impact on relative expression of FoxP3 and IL-17A. FACS analyses indicated that the addition of anti-IL-2 neutralizing antibodies had minimal effects on Sin3A sufficient cultures, while it lowered the frequency of Foxp3^+^ cells almost to significant levels in Sin3A deficient cultures. Concomitantly, neutralizing IL-2 increased the frequency of IL-17A^+^ cells only in cells lacking Sin3A, while had no effects on Sin3A sufficient controls (Fig 7D-E). Notably, the rescue in IL-17A^+^ cells was proportional to the dose of anti-IL-2 Ab (Fig 7F), independent of cell viability (Supplementary Fig. 6B), and inversely correlated with the frequency of IL-2^+^ cells (Fig 7G). Thus, inactivation of Sin3A favors IL-2 upregulation, and this appears causally linked to the upregulation of Foxp3 and the inhibition of IL-17A.

To better understand the relationship of IL-2, IL-17 and Foxp3, a logistic model was adopted for the unsupervised classification of samples based on their relative expression of these molecules. The scatter plots depicted in Fig. 8A indicate that samples were assigned to independent categories (TAM-treated and untreated) with 95% confidence only when IL-2 vs IL-17A or Foxp3 vs IL-17A were considered. Instead, when IL-2 and FOXP3 were analyzed Sin3A sufficient (untreated) and deficient (TAM treated) clustered together. Thus, together these data indicate that IL-2 producing cells are more likely to express Foxp3 rather than IL-17A. Taken together our data support a model (schematically depicted in Fig 8B) by which Sin3A normally keeps IL-2 in check over the course of Th17 differentiation balancing mTORC1, STAT3 and possibly STAT5 signaling ultimately controlling RORγt and FoxP3.

**Figure 8.**
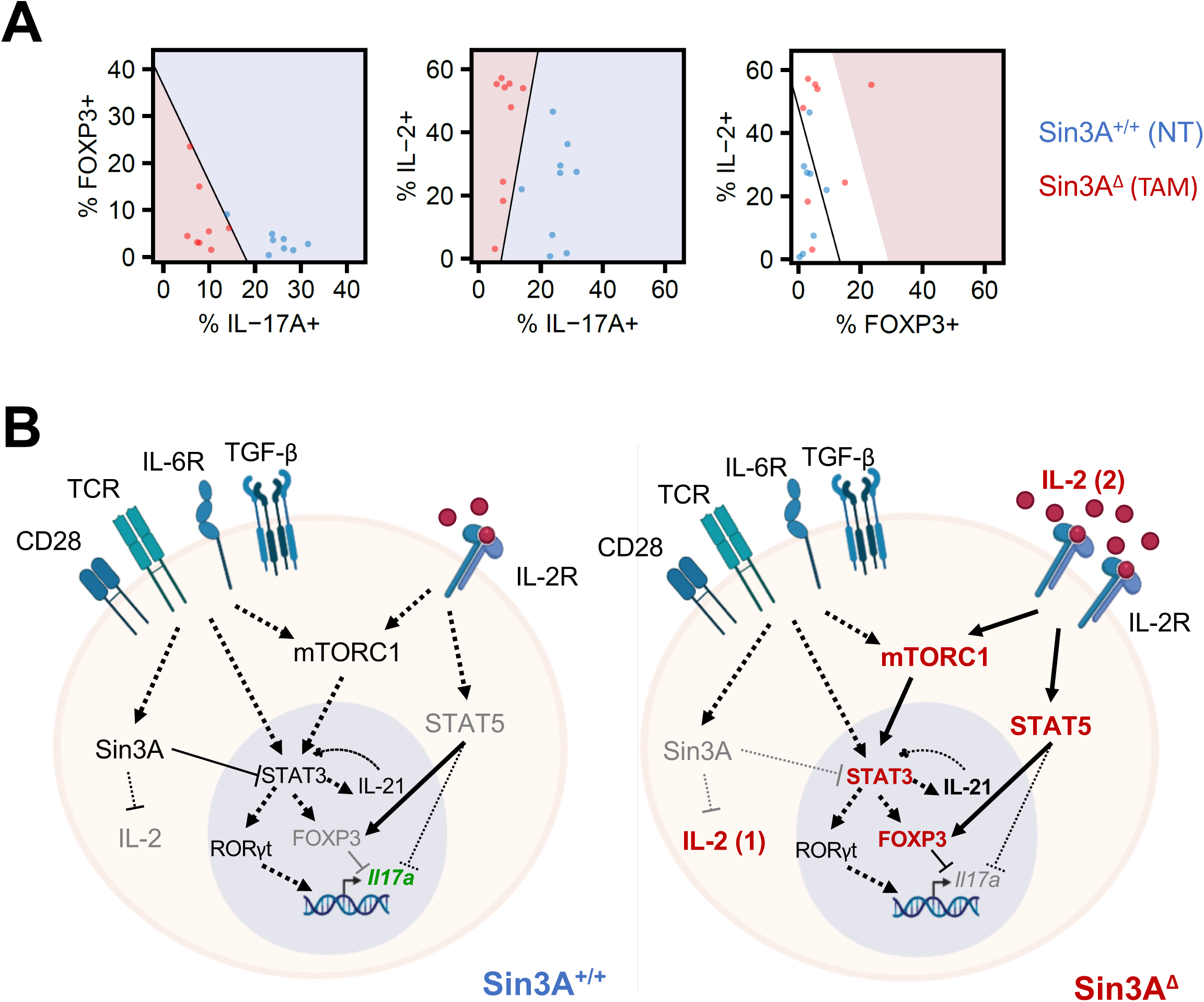
Sin3A controls Th17 cell differentiation, logistic model and schematic representation of the working model. **A.** Logistic model. The scatter plots represent the percentages of cells positive for the indicated markers. TAM-treated and untreated cells are indicated in red and blue, respectively. A logistic model was fit for the classification of samples as TAM-treated or untreated cells based on the markers shown in each plot. The red and blue areas represent the ranges in which samples are classified as TAM-treated and untreated with 95% confidence based on the fitted models, respectively. The black line represents the decision limit at which samples are equally likely to be TAM-treated or untreated. **B.** Schematic representation of data-generated working model. In Sin3A sufficient cells, Sin3A limits IL-2 expression, allowing for STAT3 and RORγt to jointly instruct the expression of *Il17a* (and of *Il17f*, *Il22* and *Il23R*). In Sin3A deficient cells, IL-2 is no longer kept under check, Foxp3 is induced, likely via mTORC1 and STAT3 signaling, hindering *Il17a* (and of *Il17f*, *Il22* and *Il23R* expression). STAT5, which activation is increased in cells lacking Sin3A, is also likely to contribute to the upregulation of Foxp3 and the inhibition of IL-17A. The figure was created with BioRender.com.

## Discussion

The transcriptional regulator Sin3A controls several pathways governing many aspects of normal and neoplastic growth and survival. Here we uncovered its critical role in CD4^+^ T cell progenitors, and over the course of Th17 differentiation.

In a previous study, Cowley *et al*. reported that Lck-Cre-driven deletion of Sin3A at the DN stage resulted in the arrest of CD8 thymocyte development, and only allowed the maturation of dysfunctional CD4SP cells(Cowley *et al*, 2005). By adopting CD4-Cre and CD4-CreER^T2^ deleter strains, we report that inactivation of Sin3A at the DP stage severely halted both CD4^+^ and CD8^+^ thymocyte maturation, causing peripheral T cell lymphopenia. Remaining CD4 T cells found in secondary lymphoid organs were found to have escaped Sin3A deletion. Thus, deletion of Sin3A at the DP stage is not compatible with further CD4 T cell development.

The reasons accounting for the different impact of Sin3A deletion, i.e., at the DN (Cowley *et al*, 2005) or DP (shown here), according to the timing remain to be fully clarified. However, it should be noted that while DN thymocytes undergo a rapid and extensive proliferative phase (6-8 cell cycle division) to ensure the generation of a large pool of progenitors, DP cells enter a resting phase to allow rearrangement of the TCR alpha chain (Taghon *et al*, 2006; Ciofani & Zúñiga-Pflücker, 2007). Indeed, by performing single cell RNA-seq analyses Li et al. found that up to 40% of intermediate SP were in the M/G1 phase, while this was not the case for DP blasts (Li *et al*, 2021). Thus, it could be speculated that deletion of Sin3A at the DN stage (Cowley *et al*, 2005) results in the selection of a cell subset capable to survive positive thymic selection, and yet lacking proper responsiveness at the mature stage. In contrast, inactivation of Sin3A at the DP stage, beyond TCR rearrangement, might render developing precursors more sensitive to negative selection. This is supported by the finding that Sin3A deficient thymocytes were enriched for apoptotic cells. A role for RORγt might also be envisaged. Indeed, it normally enforces a resting state in developing thymocytes, and favors their survival via Bcl-XL upregulation (Sun *et al*, 2000). Given the finding that in the absence of Sin3A RORγt target genes are not properly induced, whether RORγt in Sin3A deficient thymocytes retains functionality needs to be verified.

To overcome the developmental arrest and analyze the role of Sin3A in mature T cells we adopted TAM-induced deletion. We found that post-thymic Sin3A inactivation was compatible with cell proliferation and their differentiation into memory cells. Although a slight enrichment of apoptotic cells and of cells arrested in the S-phase in cultures was detected within Sin3A-deficient cells, previously reported in other model systems by us and others (Dannenberg *et al*, 2005; McDonel *et al*, 2012; Gambi *et al*, 2019), Sin3A sufficient and deficient (TAM-treated) cells completed comparable rounds of cell division and acquired several memory markers. By day 3, Sin3A knock down cells also revealed a distinct mTORC1 signaling signature, corroborated by higher CD25 and PD-1 surface levels, and increased phosphorylation of p70S6K, an mTORC1 effector molecule (Brunn *et al*, 1996). Nevertheless, in Th17 skewing conditions, Sin3A deficient cells failed to produce IL-17A because of *Il17a* defective expression. Compared to controls, Sin3A deficient T cells also failed to upregulate *Il17f*, *Il23* and *Il22*, typical determinants of Th17 cell differentiation, and instead upregulated *Il21* to higher extents when compared to controls. As *Il21* is more dependent on STAT3 (Wei *et al*, 2007), while *Il17a*, *Il23R* and *Il22* require RORγt (Zhou *et al*, 2007), the gene expression profile of Sin3A deficient cells was consistent with the cells having STAT3 (increased phosphorylation and nuclear representation), but RORγt transcriptionally active.

Our data support the possibility that RORγt malfunctioning was related to the acquisition of Foxp3. Indeed, we found Sin3A deficient cells to express either Foxp3 or IL-17A by a mutually exclusive modality, suggesting the existence of a causative link. In line with this possibility, we found that the Foxp3 P60 inhibitor, able to bind to Foxp3 and inhibit its nuclear translocation and activity (Casares *et al*, 2010), increased the percentage of IL-17A-expressing cells most significantly in Sin3A deficient cells. Previously published studies support our findings. Zhou and co-authors (Zhou *et al*, 2008) had found that TGFβ-induced Foxp3 could inhibit RORγt through a direct interaction, also involving Foxp3 DNA binding domain (Zhou *et al*, 2008). In their study these authors reported that RORγt and Foxp3 were co-expressed in naive CD4^+^ T cells exposed to TGF-β, and that IL-6, IL-21 and IL-23 were sufficient to relieve Foxp3-mediated inhibition of RORγt, thereby promoting Th17 cell differentiation.

Foxp3 expression in human and mouse T cells is induced by STAT3 activation downstream to CD3/CD28 and TGFβ (Pallandre *et al*, 2007). Also, IL-2 promotes Foxp3 expression by increasing its stability (Chen *et al*, 2011). We found that Sin3A deficient T cells expressed higher levels of IL-2, and that neutralizing anti-IL-2 antibodies limited Foxp3 upregulation and partially rescued IL-17A expression in Sin3A deficient cells. Lam et al. previously found that Sin3A represses IL-2 gene expression by bringing histone deacetylase activity to the IL-2 promoter. Data indicated that TCR-induced Cdk5-dependent Sin3A phosphorylation and destabilization, reduces Sin3A/HDAC binding to the IL-2 promoter, allowing higher IL-2 expression (Lam *et al*, 2015). Our data are consistent with this previous report, and with the possibility that Sin3A negatively regulates IL-2 expression in Th17 conditions to restrain Foxp3 induction and enable RORγt activity.

Other mechanisms might further contribute to the observed phenotype. Indeed, blocking Foxp3 or neutralizing IL-2 only partially rescued IL-17-producing cells. For instance, Sin3A through the binding to p300 and HDAC1 (Tsai *et al*, 2017; Kadamb *et al*, 2013; Dancy & Cole, 2015) might control RORγt (Wu *et al*, 2015) and/or Foxp3 (Liu *et al*, 2013, 2014; Xiao *et al*, 2014) acetylation, and by that their activities. In addition, a number of transcription factors fine tune Th17 differentiation (Tanaka *et al*, 2014; Ciofani *et al*, 2012; Zhang *et al*, 2008; Yang *et al*, 2007b). We found that STAT5 phosphorylation was increased in cells lacking Sin3A. Given its ability to favor Foxp3 expression (Burchill *et al*, 2007) and to inhibit Th17 cell differentiation (Laurence *et al*, 2007), also STAT5 could contribute to the loss of IL-17A^+^ cells and the appearance of the Foxp3^+^ ones. We are in the process of further interrogating the RNA-seq data to uncover whether additional mechanisms are indeed contributing to the phenotype.

To investigate the effect of Sin3A inactivation in vivo, we attempted the transfer of finite number of T cells in syngenic recipients and treated them with TAM. Sin3A deletion was observed, and yet cells failed to persist (not shown). So, whether events similar to those described here shape Th17 cell function *in vivo* remains to be determined. Nevertheless a subset of T cells within the small intestinal lamina propria was found to co-express both Foxp3 and RORγt and to produce less IL-17 than those that express RORγt alone (Zhou *et al*, 2008), suggesting that Th17 cells can acquire distinct physiological functions in response to extracellular signals (cytokines/growth factors). We suggest that Sin3A contributes to such differentiation program, by tuning IL-2 and mTORC1 and ultimately Foxp3. Although some controversy regarding the role of mTORC1 signaling on Foxp3 expression exists (Battaglia *et al*, 2005; Heather Kopf, Gonzalo M. de la Rosa, O.M.Zack Howard, 2007), it has been shown to regulate the generation and function of central and effector Foxp3^+^ regulatory T cells (Sun *et al*, 2018). It is thus tempting to suggest that IL-2 and/or agents directly tuning mTORC1 (i.e., nutrients, microbial components) (McGeachy & McSorley, 2012) might shape the inflammatory/regulatory phenotype of Th17 cells. As Sin3A inhibitors have been developed for *in vivo* use, these data raise the possibility to attempt the inhibition of Sin3A (Kwon *et al*, 2015) in autoimmune diseases or gut inflammatory disorders. In these conditions, Sin3A inhibition could dampen the pro-inflammatory nature of Th17 cells, while favoring immunoregulatory functions.

## Acknowledgements

The authors acknowledge funding from Associazione Italiana sulla Ricerca sul Cancro (AIRC: IG 2018 Id.21763), and from a generous anonymous donor. The authors are grateful to Dr. Jan Tavernier (Ghent University) for the provision of Sin3A^F/F^ mice, Dr. Paolo Dellabona (San Raffaele Institute) for the provision of CD4-Cre mice, and Dr. Marco Bianchi (San Raffaele Institute) for the provision of Rosa26-ER^T2^Cre mice. Finally, the authors would like to thank Dr. Maya Fedeli (San Raffaele Institute) for help with thymic analysis and for anti-IL-2 antibodies, Dr. Christopher Bruhn (FIRC Institute of Molecular Oncology, IFOM, Milan, Italy) for help with the logistic models and present and past members of the Mondino’s lab for frequent discussion.

## Author contributions

LP performed experiments, analyzed data and prepared the manuscript; LI initiated the project, performed proof of concept experiments and backcrossed mice; EDS and VB performed experiments; AA and FPC performed RNA seq; TL and JJL analyzed RNA seq data, provided the p60 inhibitor and critical suggestions; AM supervised activities and contributed to experimental design, data analyses and interpretation and manuscript preparation.

