## Supplemental figures and legends for "The transcriptional regulator Sin3A balances IL-17A and Foxp3 expression in primary CD4 T cells"

**Supplementary figure legends**

**Supplementary Figure 1. Tamoxifen-induced Sin3A deletion *in vivo* hinders CD4 T cell peripheral representation. A.** Experimental scheme. **B.** Sin3A$^{F/F}$ CD4-CreERT2$^{+/-}$ and Sin3A$^{F/F}$ CD4-CreERT2$^{-/-}$ were treated with Tamoxifen for 5 consecutive days. CD4 T cell representation in the blood was monitored overtime. Individual mice are depicted.

**Supplementary Figure 2. Sin3A inactivation impacts on T cell survival.** Sin3A$^{F/F}$ Rosa26-CreER$^{T2}$ CD4$^{+}$ T cells were treated as described in Figure 2. **A.** Annexin V/7-AAD staining and **B.** quantification N=4, **C**. Cell cycle analysis. Error bars: SD. Two-tailed paired Student's T-test.

**Supplementary Figure 3: Memory phenotypes of Sin3A sufficient and deficient cells kept in Th17 skewing conditions**. Flow cytometric analyses of CD44, CD62L, CD25, PD-1 and CD69 in untreated (NT) and treated (TAM) Sin3A$^{F/F}$ Rosa26-CreER$^{T2}$ CD4$^{+}$ T cells **A.** cultured in IL-7 or **B.** Th17 conditions. **C.** Relative expression (RFI, relative fluorescence intensity) of CD44, CD69, CD25 and PD-1 in untreated (NT) and treated (TAM) Sin3A$^{F/F}$ Rosa26-CreER$^{T2}$ CD4$^{+}$ T cells in Th17 conditions.

**Supplementary Figure 4: Antigen-driven activation of Sin3A sufficient and deficient cells**. **A.** Proliferation profiles of Sin3A$^{+/F}$ CreERT2$^{+/-}$ OTII$^{+/-}$ and Sin3A$^{F/F}$ CreERT2$^{+/-}$ OTII$^{+/-}$ T cells untreated (NT) or treated with TAM over a 3 days ovalbumin-driven culture. **B.** quantification of A. **C.** Percentages of live, CD25+ and CD44+ cells. N=3. Error bars: SD. Two-tailed paired Student's T-test.

**Supplementary Figure 5: Defective IL-17A expression in Sin3A-deficient cells is observed in cells with comparable proliferation status.** Representative dot plots and of Th17-skewed Sin3A$^{F/F}$ Rosa26-CreER$^{T2}$ CD4+ T cells, treated with 4-OH tamoxifen (TAM) or left untreated (NT).

**Supplementary Figure 6: Cell viability is preserved in spite of Foxp3 or IL-2 inhibition.** Cells were cultured as depicted in Figure 2 in the absence or the presence of the P60 Foxp3 inhibitor **(A)** or a neutralizing anti-IL-2 mAb (**B**). The fraction of viable cells in the various culture conditions is shown. Individual experiments are depicted. Error bars: SD. Two-tailed paired Student's T-test. N=5 (A), N=10 (B).

**Supplementary Figure 7. STAT5 and p70S6K phosphorylation levels are increased in cells lacking Sin3A.** Sin3A$^{F/F}$ Rosa26-CreER$^{T2}$ CD4$^{+}$ T cells were left untreated or treated with TAM as depicted in Figure 2. Representative histograms (**A, C**) and relative expression in independent experiments (**B, D**) are depicted. N=3. Error bars: SD. One-tailed paired Student's T-test.

| Primer | | Sequence 5'-3' |
|---|---|---|
| Il17a | Fw | CAGGGAGAGCTTCATCTGTGT |
| | Rv | GCTGAGCTTTGAGGGATGAT |
| Ifng | Fw | GCGTCATTGAATCACACCTG |
| | Rv | ATCAGCAGCGACTCCTTTTC |
| Il17f | Fw | CCCAGGAAGACATACTTAGAAGAAA |
| | Rv | CAACAGTAGCAAAGACTTGACCAT |
| Rorc | Fw | ACCTCTTTTCACGGGAGGA |
| | Rv | TCCCACATCTCCCACATTG |
| Rora | Fw | AACCCGAACCCATATGTGAC |
| | Rv | ATGTTCTGGGCAAGGTGTTC |
| Il21 | Fw | AGGAGGGGAGGAAAGAAACA |
| | Rv | GGGAATCTTCTCGGATCCTC |
| Il23r | Fw | AAGGCTTTTCGGAACCTCAT |
| | Rv | TTCCAGGTGCATGTCATGTT |
| Foxp3 | Fw | AGAAGCTGGGAGCTATGCAG |
| | Rv | GCTACGATGCAGCAAGAGC |
| Il2 | Fw | GCTGTTGATGGACCTACAGGA |
| | Rv | TTCAATTCTGTGGCCTGCTT |
| UbC | Fw | TGGCTATTAATTATTCGGTCTGCA |
| | Rv | GCAAGTGGCTAGAGTGCAGAGTAA |
| Il22 | Fw | TCGCCTTGATCTCTCCACTC |
| | Rv | GCTCAGCTCCTGTCACATCA |

Supplementary table 1

| **Antibody** | **Fluorophore** | **Company** | **Cat #/clone** |
|---|---|---|---|
| CD5 | APC/PE | Biolegend/BD | Clone 53.7.3 |
| TCRβ | PB | Biolegend | 109226 |
| TCRβ | FITC | BD | Clone H57-597 |
| CD3 | APCCy7 | Biolegend | Clone 145-2C11 |
| CD8 | Diverse | Biolegend/BD | Clone 53-6.7 |
| CD4 | Diverse | Biolegend/BD | Clone RM4-5 |
| CD44 | PCPCy5.5 | Biolegend | 103032 |
| CD44 | PE | BD | 553134 |
| CD62L | APC | BD | 553152 |
| CD62L | PECy7 | BD | 560516 |
| IL-17A | APCCy7 | Biolegend | 506940 |
| IFNγ | PE | Biolegend | 505808 |
| RORγt | APC | eBioscience | 17-6988-82 |
| IL-2 | PECy7 | Biolegend | 503832 |
| FOXP3 | PE | eBioscience | 12-5773-82 |
| PD1 | PECy7 | Biolegend | 135216 |
| CD25 | PE | BD | 553866 |
| CD69 | APCCy7 | Biolegend | 104525 |
| AnnexinV | APC | BD | 550474 |
| Sin3A | none | Abcam | ab129087 |
| Rb IgG isotype control | none | Abcam | ab172730 |
| Rb IgG | FITC | BD | 554020 |
| pSTAT5 | none | Cell Signaling | 9351 |
| pP70-S6K (T389) | none | Cell Signaling | 9205 |

Supplementary table 2

A

B

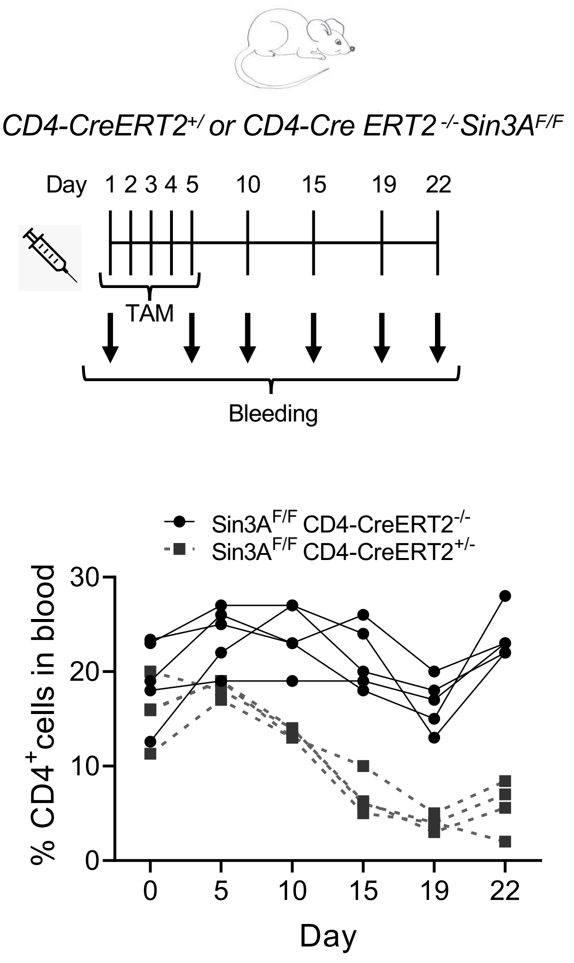


Supplementary figure 1

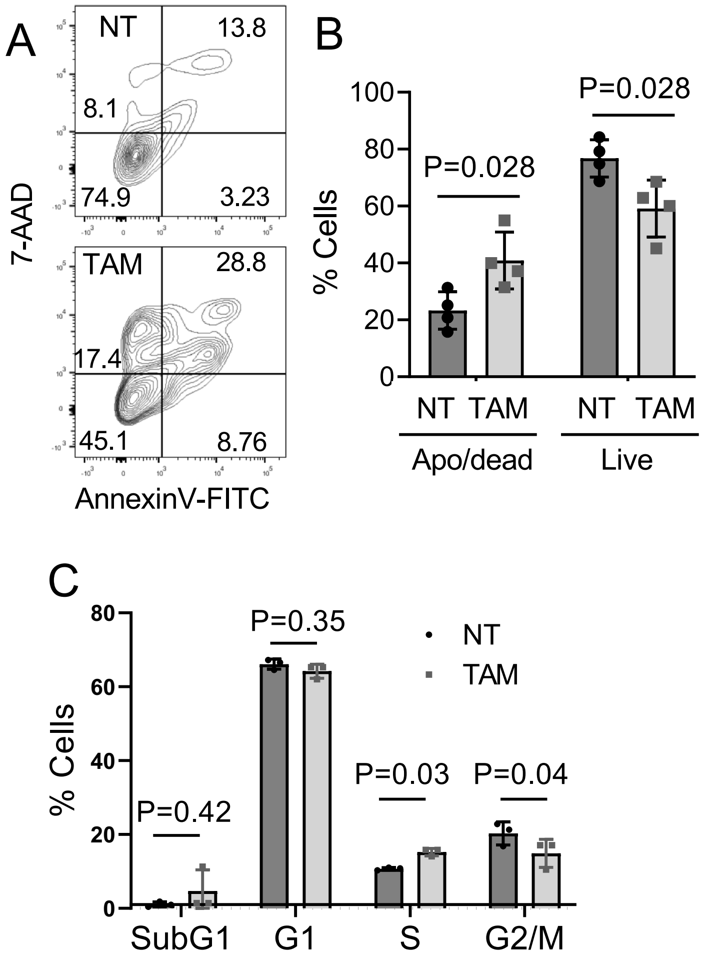


Supplementary figure 2

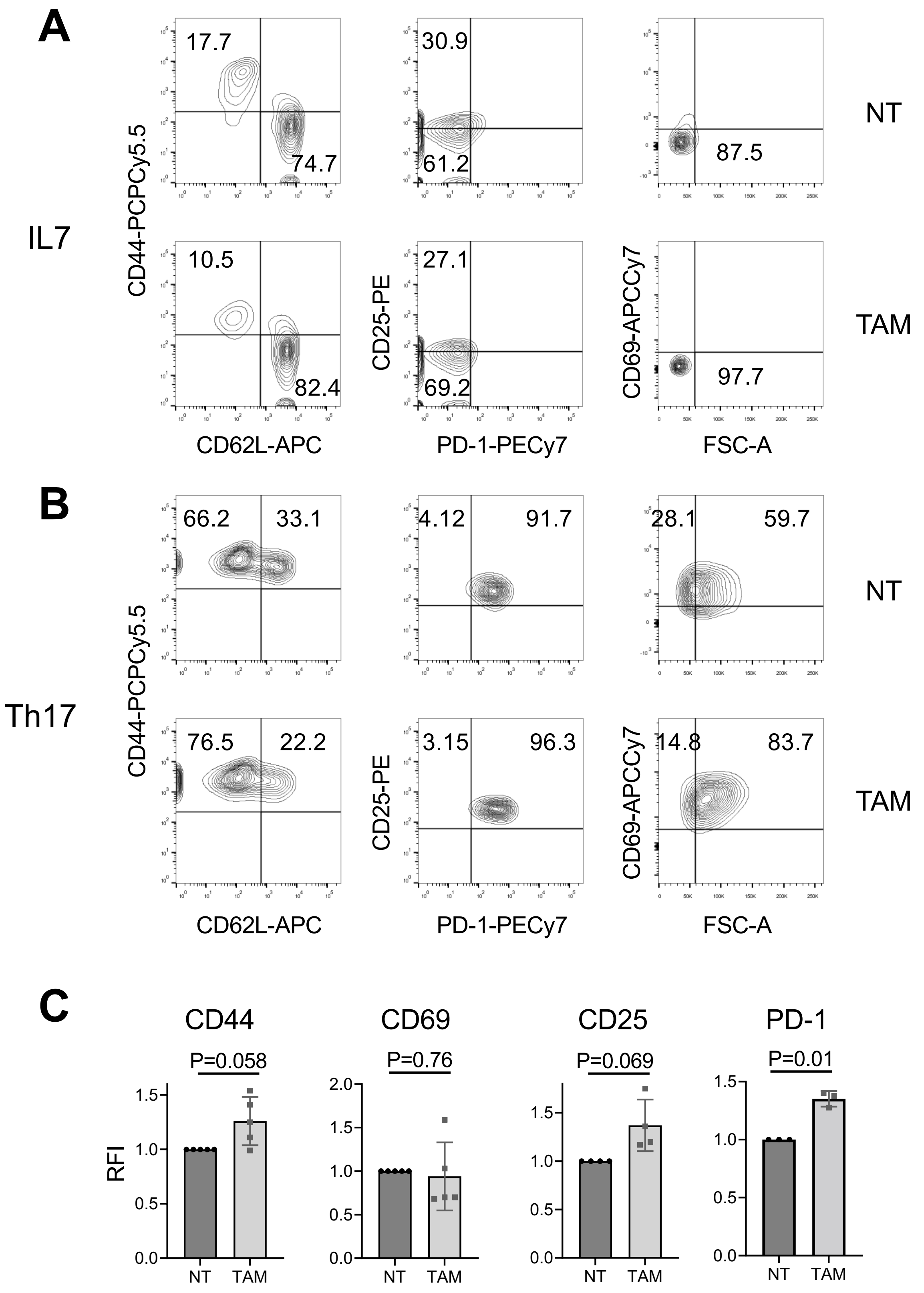


Supplementary figure 3

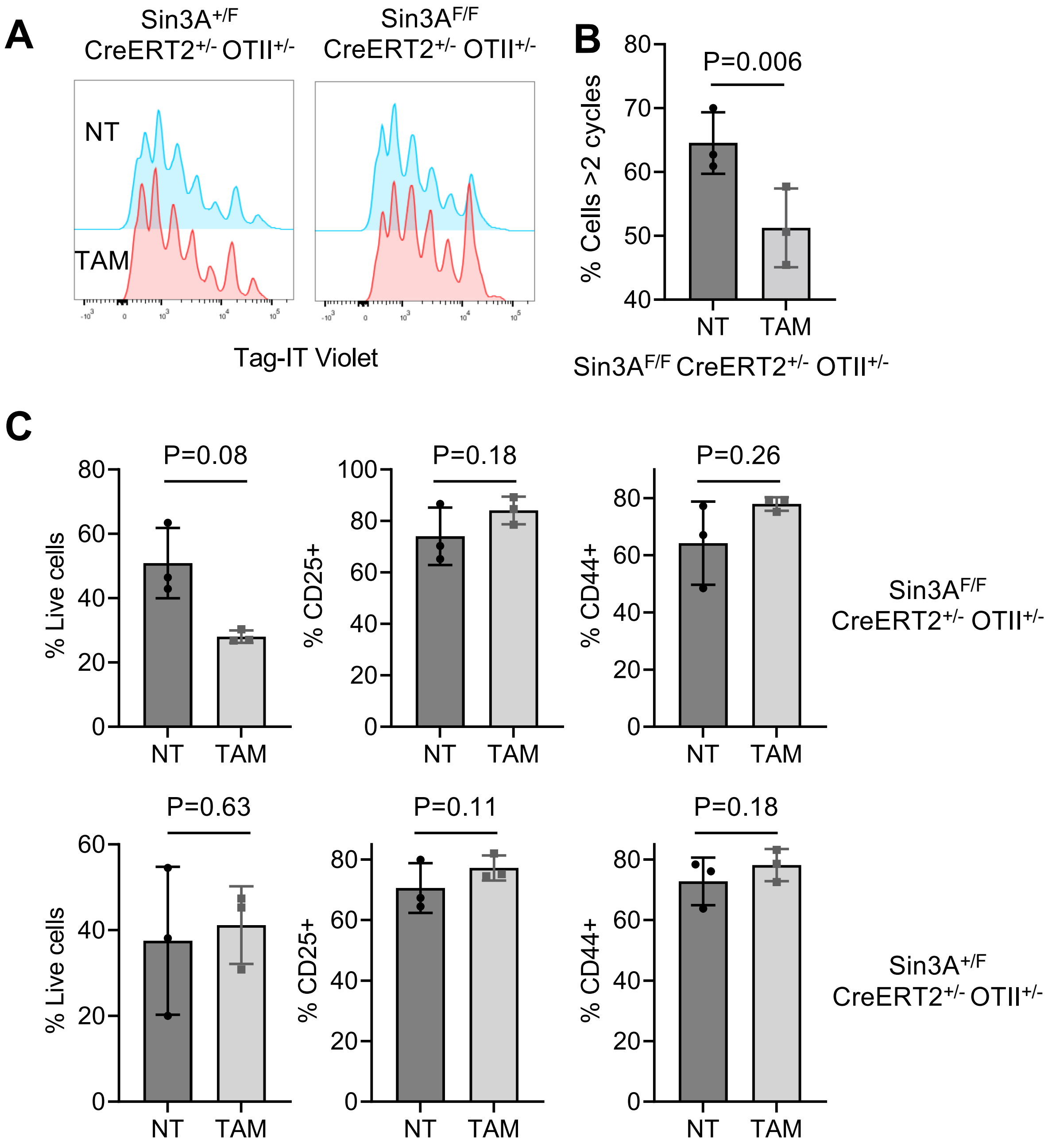


Supplementary figure 4

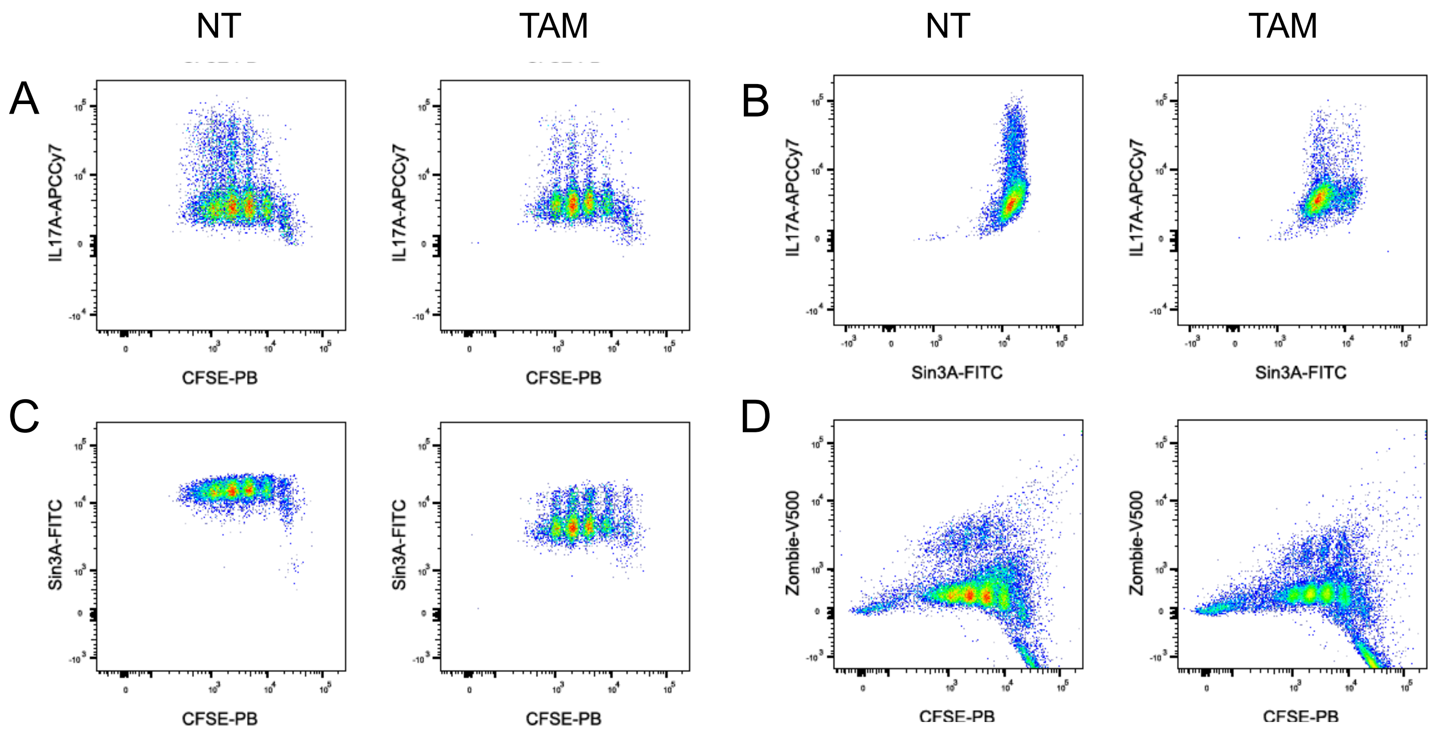


Supplementary figure 5

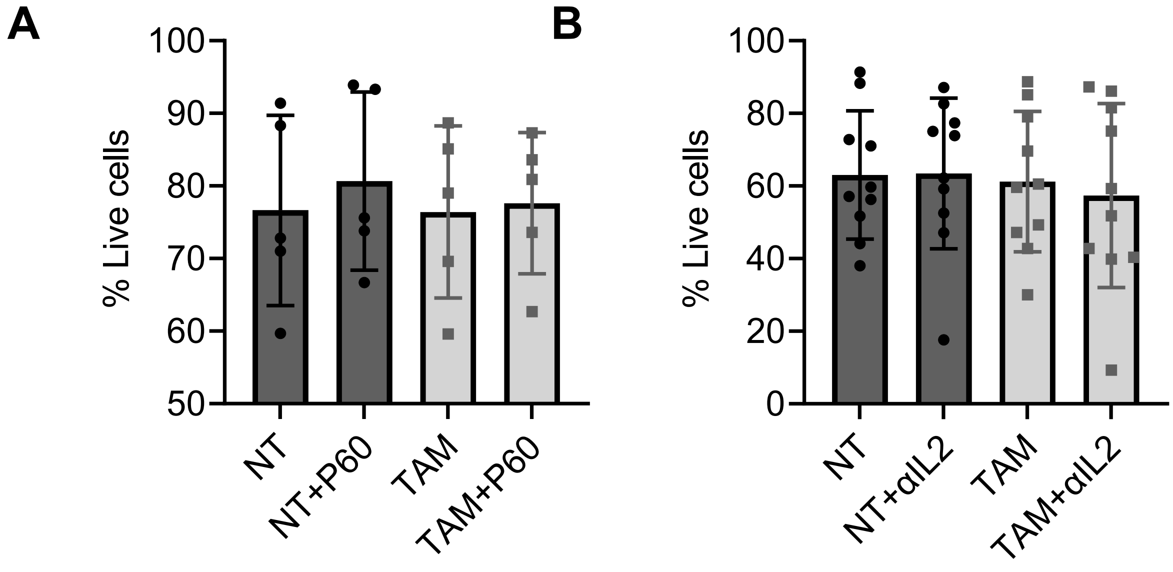


Supplementary figure 6

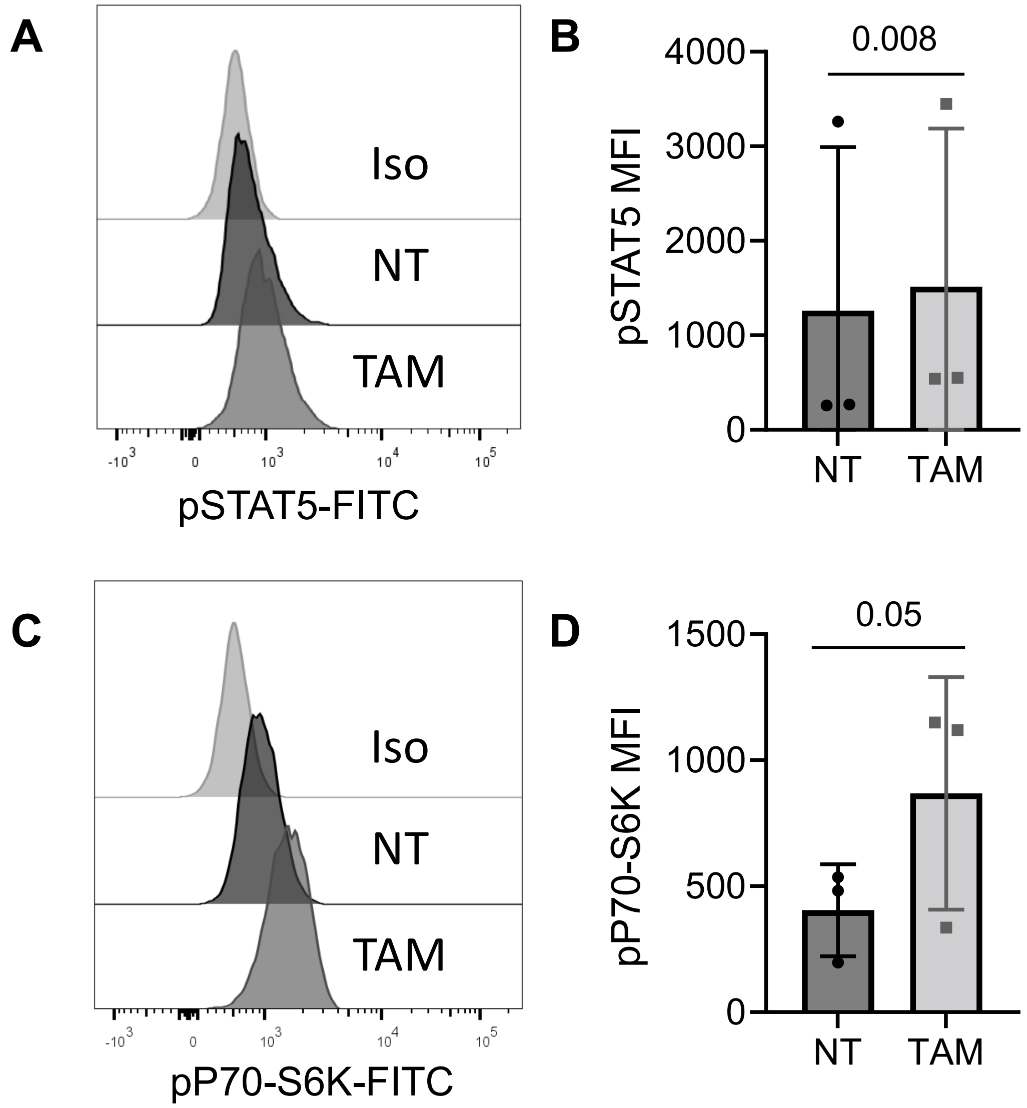


Supplementary figure 7
